# Scaling *k*-Means for Multi-Million Frames: A Stratified NANI Approach for Large-Scale MD Simulations

**DOI:** 10.1101/2025.06.15.659780

**Authors:** Jherome Brylle Woody Santos, Lexin Chen, Ramón Alain Miranda-Quintana

## Abstract

We present improved *k-*means clustering initialization strategies for molecular dynamics (MD) simulations, implemented as part of the *N*-ary Natural Initiation (NANI) method. Two new deterministic seeding strategies: strat_all and strat_reduced, extend the original NANI approaches and dramatically reduce the clustering runtime while preserving the quality of clustering results. These methods also preserve NANI’s reproducible partitioning of well-separated and compact clusters while avoiding the costly iterative seed selection procedures of previous implementations. Testing on the β-heptapeptide and the HP35 systems shows that these new flavors achieved Calinski-Harabasz and Davies-Bouldin scores comparable to the previous NANI variant, indicating that the efficiency gains come with no quality decrease. We also show how these new variants can be used to greatly speed up our previously proposed Hierarchical Extended Linkage Method (HELM). These enhancements extend the reach of NANI to accelerate large-scale MD analysis both in stand-alone *k*-means clustering and as a component of hybrid workflows, and remove a key barrier to routine, scalable, and reproducible exploration of complex conformational ensembles. The improved NANI implementation is accessible through our MDANCE package: https://github.com/mqcomplab/MDANCE.

## 1. INTRODUCTION

Clustering has become an important tool in molecular dynamics (MD) trajectory analysis. It is widely used to probe structure, dynamics, and energetics of biomolecular systems across time.^1–3^ Among various clustering algorithms, *k-*means remains one of the most widely used due to its computational efficiency.^4^ However, conventional *k-*means has some well-known drawbacks for its applications on MD data. In particular, the algorithm is highly sensitive to the initial centroid selection, and the common seeding schemes (random or *k-*means++ initialization) can perform poorly on complex and high-dimensional MD datasets, while also having great variability in the final results. *k*-means++ uses a probability based greedy initialization that improves upon random seeding, but being stochastic, it still cannot guarantee reproducible results.^5,6^ This lack of reproducibility is also a serious issue for MD analysis as different runs can lead to different cluster assignments, which may hinder the extraction of meaningful physical insights from these trajectories.

To address these challenges, the *k-*means *N*-ary Natural Initiation (NANI) protocol was recently developed.^7^ NANI introduces a new seeding approach by using efficient *n*-ary comparisons, which involve evaluating multiple objects simultaneously,^8–15^ to identify high-density regions of the conformational space and select diverse, representative initial seeds. By design, the selected frames are both dense and well-separated, allowing the *k-*means algorithm to generate well-partitioned clusters. More importantly, NANI is completely deterministic: it produces identical clusters and populations across repeated runs, removing stochastic variability and enhancing reproducibility. Previous benchmarks show that NANI clusters are compact, well-separated, and in agreement with known conformational states.

In its initial implementation, NANI provided two deterministic seeding modes: comp_sim and div_select. The former selects initial centers based on the diversity within the densest region of the dataset, while the latter maximizes diversity across the entire dataset. Both strategies improved clustering quality and reproducibility compared to random or *k-*means++ seeding. However, as the size of MD datasets continues to increase, even these strategies encounter performance bottlenecks due to the high computational cost of processing massive numbers of frames. In this approach, we found that the main computational bottleneck is the diversity selection algorithm, which can scale poorly to millions of frames.

In this manuscript, we introduce two new seeding strategies for NANI: strat_all and strat_reduced. These methods preserve NANI’s deterministic character and robustness, while substantially accelerating the overall seeding process. We outline these stratified initialization approaches and demonstrate their performance on two representative MD systems: β-heptapeptide and villin headpiece (HP35). We use standard validation metrics like the Calinski-Harabasz index (CHI, also known as Pseudo-F score)^16^ and Davies-Bouldin index (DBI)^17^ to evaluate cluster compactness, separation, and consistency across the initialization schemes. The results show that strat_all and strat_reduced match the clustering performance of original NANI methods, while significantly reducing the computational time. A single *k*-means pass is ∼1.5x faster for the β-heptapeptide, ∼13x for 150k HP35 frames, and nearly 45x for 1.5 M HP35 frames: speedups that directly translate to faster HELM clustering with no loss of accuracy.

## 2. THEORY

### 2.1. *k*-means initialization and overview of NANI

*k*-means clustering requires an initial set of *k* cluster centers (seeds), which are traditionally chosen at random or via *k-*means++. NANI provides a deterministic alternative: it ranks frames by how representative they are of the ensemble and then selects seeds that are both high-density (representative) and well-separated (diverse). This reduces run-to-run variability and improves reproducibility.

### 2.2. Complementary similarity and cMSD

NANI uses complementary similarity, a concept that follows *n*-ary similarity to measure how much the overall similarity of the dataset changes when a single frame/conformation is removed. We report this as the complementary mean squared deviation (cMSD): frames with high cMSD are those whose removal decreases the dataset’s overall similarity the most. In practice, high-cMSD frames tend to be the central/representative conformations (which we consider as the set’s medoid), making cMSD an efficient alternative for assessing structural density.

### 2.3. Original seeding strategies

Early NANI implementations introduced two deterministic seed-selection modes: comp_sim and div_select. In comp_sim, we (i) compute cMSD for all frames, (ii) retain only a user-specified top fraction (e.g., top 10%) as pool of high-density candidates, and then (iii) select seeds iteratively by choosing the frame that is “farthest” from the already selected seeds (starting from the medoid). div_select uses the same iterative diversity maximization but applies it to the full dataset without the initial density filter. Both strategies produce diverse, representative seeds, but the iterative selection, which requires the repeated scanning of the remaining candidates, can be costly as the dataset size grows.

### 2.4. New stratified seeding strategies

To avoid the iterative distance maximization, the latest NANI version introduces two single-pass, deterministic strategies: strat_all and strat_reduced. Both rely on cMSD to sort frames from most to least representative, then selects seeds by evenly spaced ranks from this ordered list. Intuitively, these methods sample across the full range of high-density conformations while avoiding the repeated candidate-to-seed distance scans used by the earlier methods (Fig. 1).

**Figure 1.**
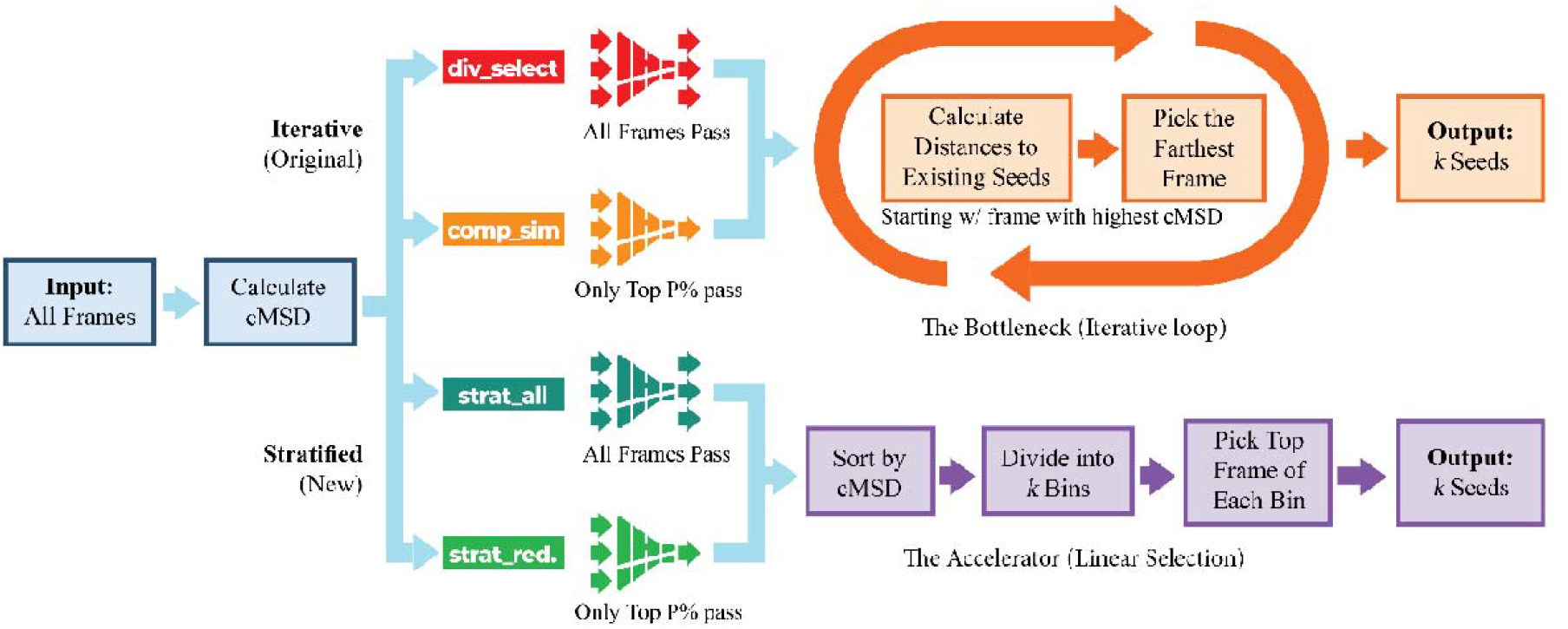
Schematic diagram of the four NANI seeding strategies. All methods begin by computing per-frame cMSD. div_select and comp_sim then select seeds iteratively by repeatedly choosing the frame farthest from the current seed set (with comp_sim restricting candidates to a top-cMSD subset). strat_all and strat_reduced replace iterative selection with a single-pass stratified procedure: frames are ranked by cMSD, divided into bins, and the top frame from each bin is used to form the candidate seed list (for strat_reduced, binning is applied only within the top-cMSD subset). The output in all cases is a deterministic set of *k* initial seeds for *k*-means clustering.

#### strat_all

In strat_all, we perform stratified sampling over the entire dataset. First, we compute the cMSD for all frames and sort them from highest to lowest. A user-defined percentage parameter *p* determines how many candidates are selected from this ranked list. Unless otherwise stated, we use *p* = 10%; sensitivity of clustering quality to *p* is reported in the Supporting Information. If the dataset has *N* frames, we select

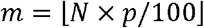

frames by dividing the list into *m* equal-sized strata (bins). From each bin, the top-ranked frame (highest cMSD) is selected as a representative. The first *k* of these selected frames are then used as the initial *k*-means seeds. With this procedure, strat_all ensures that the seeds include the most important conformations while also covering different regions of the space. Furthermore, there are no random steps made, given the same dataset and percentage parameter, strat_all will always produce the same set of seeds. The complexity is primarily in the calculation and sorting of the cMSD values, which are far more efficient than an exhaustive iterative selection.

#### strat_reduced

For strat_reduced, we follow the same stratification idea but apply it only within a high-density subset. First, we identify the top fraction of frames with the highest cMSD (as in comp_sim). Within this subset, we recompute cMSD and apply the same stratified selection procedure used in strat_all to obtain a diverse list of candidate seeds; the first *k* are used as the initial centers. This approach effectively filters out less representative outliers before selecting seeds. As a result, strat_reduced focuses initialization on dense regions while still spreading initial centers to improve diversity.

### 2.5. Efficiency and scalability

Both strat_all and strat_reduced eliminate the iterative seed-addition loop with a one-time selection from a ranked list of indices. Their dominant costs are on computing cMSD (O(*N*)) and sorting O(*N*·log *N*). By contrast, comp_sim and div_select repeatedly evaluate distances as seeds are added, which can scale poorly for large *N*. In practice, the stratified methods substantially reduce initialization time at MD scale while preserving determinism.

### 2.6. Baseline and comparisons

Extensive comparisons between the original NANI methods and conventional initialization strategies (e.g., random, *k*-means++) have already been reported in our prior work,^7^ where NANI was shown to outperform these methods in both clustering quality and reproducibility. Those analyses have also demonstrated that the comp_sim mode is significantly more computationally efficient and more robust than div_select. Therefore, in this work we use comp_sim as the baseline and focus comparisons on the two new stratified strategies (strat_all and strat_reduced).

## 3. SYSTEMS

### β-Heptapeptide

The topology and trajectory files were obtained from publicly available data and accessed via GitHub.^18,19^ Clustering was performed using only the backbone atoms (N, Cα, C, O, H) of residues 2 to 11; excluding terminal residues and all side chains to reduce noise, consistent with the protocol described by Daura et al.^20^ After discarding the first 1,000 frames as equilibration, the trajectory was aligned to the first frame. A total of 6,001 structures were used for clustering.

### HP35

We analyzed the long-timescale trajectory of the Nle/Nle mutant of the C-terminal sub-domain of the villin headpiece (HP35) from D. E. Shaw Research.^21^ The production run spans 305 μs at 360 K and contains ∼1.52 million frames, saved every 200 ps. Frames before 5,000 were discarded as an equilibration period, and the remaining frames were aligned to the 5000^th^ frame. Clustering was performed using the backbone heavy atoms, following conventions established in previous HP35 studies: the N atom of residue 1; the N, Cα, and C atoms of residues 2 to 34; and the N atom of residue 35. Sieving was applied to select every 10th frame, yielding a more manageable subset of 151,104 structures. Clustering was performed on both this sieved subset (used for the main clustering and HELM analyses) and the full 1.5 million-frame trajectory (used for runtime scaling benchmarks).

The trajectory-level conformational variability for both systems is summarized in the Supporting Information using RMSD-to-first frame traces and distributions (Figs. S6-S9).

## 4. RESULTS & DISCUSSION

### 4.1. Clustering results and validity metrics

We evaluated the three NANI variants (comp_sim, strat_all, and strat_reduced) on the β-heptapeptide and the HP35 systems. Our initial test involves the screening on the number of clusters from 5 to 30: for each *k* we ran *k-*means with the calculated seeds and computed two standard internal validation metrics: Calinski-Harabasz (CHI) and Davies-Bouldin (DBI) indices. The CHI describes the ratio of the inter and intra-cluster dispersion; larger values signify tighter, better-separated clusters.^16^ On the other hand, the DBI quantifies how well the clusters are both compact and mutually separated; lower values indicate greater inter-cluster separation and intra-cluster compactness.^17^

As shown in Fig. 2, every seeding strategy within NANI yields nearly overlapping CHI curves on both systems. While the DBI curves show visual differences, they follow the same overall trend for all three initiators. These observations confirm that across the full screening window, the stratified NANI variants deliver clustering performance that matches the quality of the original NANI method.

**Figure 2.**
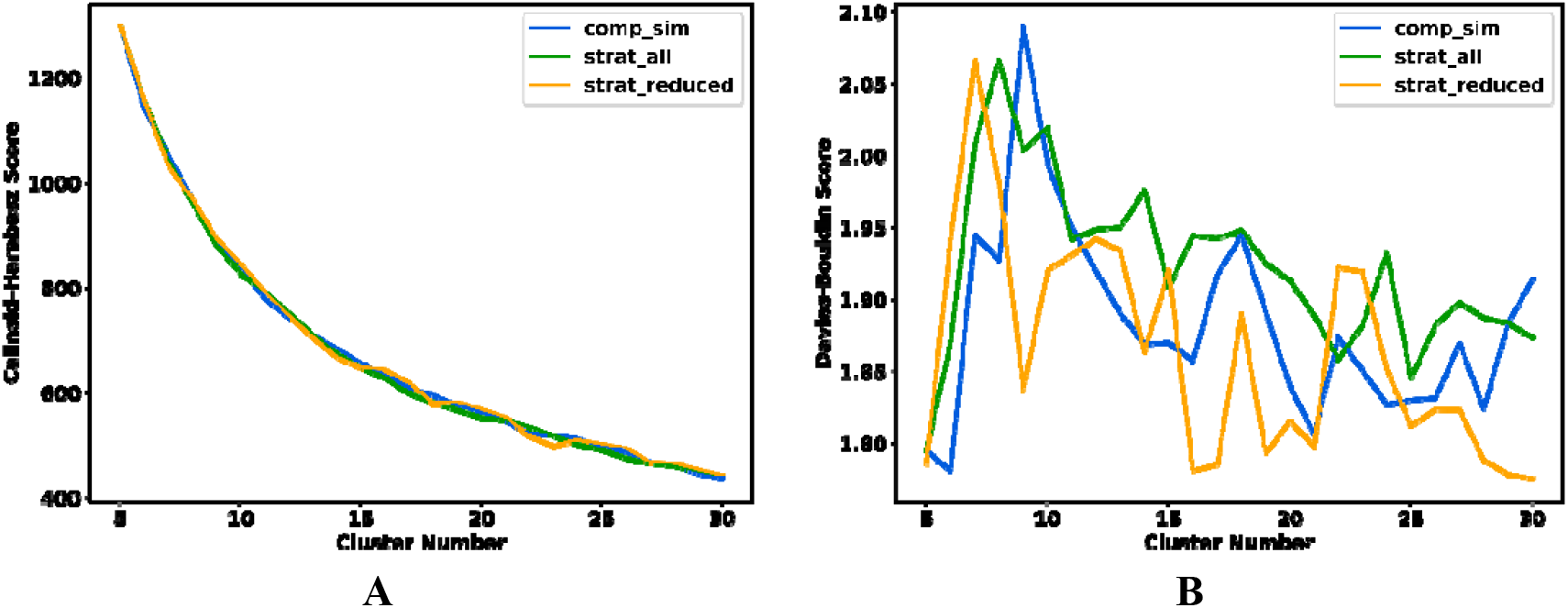
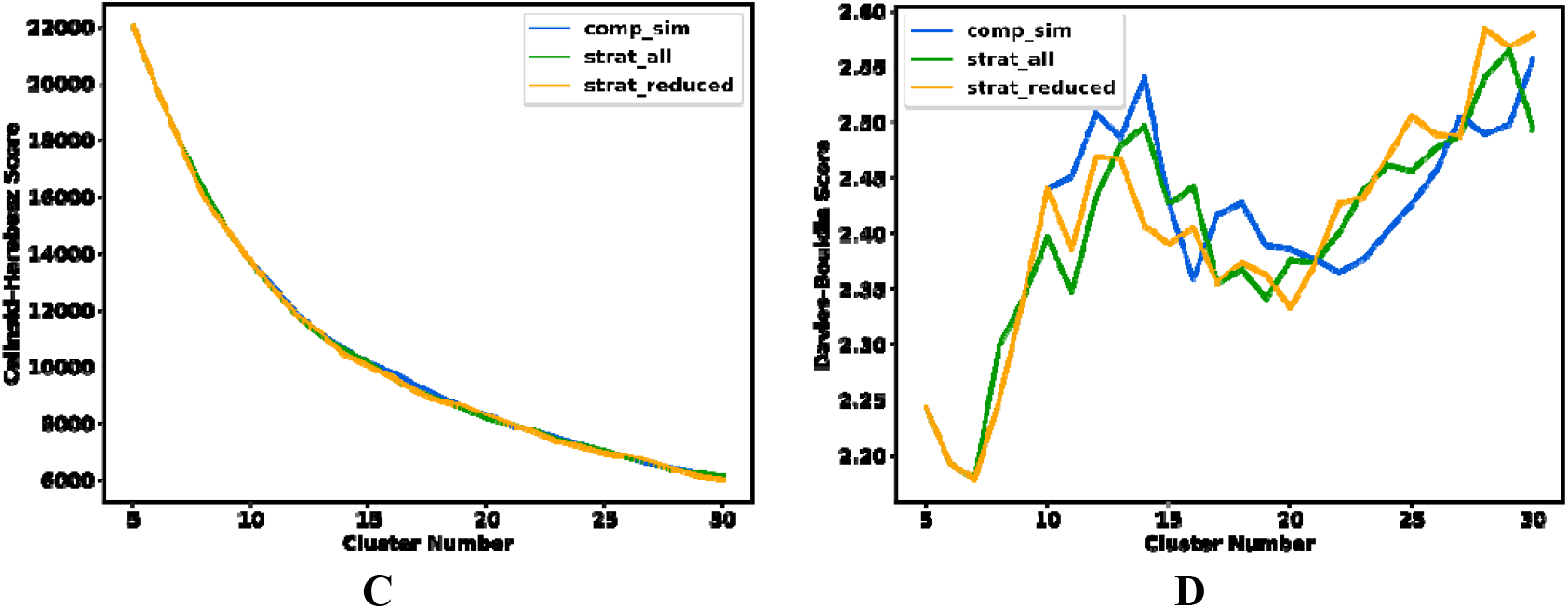
Change in Calinski-Harabasz (A, C) and Davies-Bouldin (B, D) indices for (A, B) β-heptapeptide and (C, D) HP35 after *k-*means NANI screening from *k* = 5 to *k* = 30.

For the stratified strategies, we also assessed the sensitivity to the percentage parameter *p* (5% to 30%). Over this range, the DBI trends remain consistent (Fig. S5).

As we have shown in previous studies, the CHI and DBI curves are valuable not only for comparing clustering methods (or initialization strategies in this case), but also for estimating the most meaningful cluster count for a given trajectory. We approach this in two ways. First, a global assessment of the curves or simply the selection of the absolute CHI maximum or DBI minimum across the screened range. Second, a local stability assessment of the curvatures of the metrics by evaluating their finite-difference second derivatives, the prominent extrema in these curvatures might be useful in identifying *k* values that are more stable than their immediate neighbors. As both indices are known to favor small cluster counts, the search was limited to *k* ≥ 5.

In Table 1, we summarize the cluster counts suggested by global criteria (absolute DBI minimum and absolute CHI maximum) for each seeding strategy. For the β-heptapeptide, the CHI is fully consistent across methods (*k* = 5 to 6), whereas the DBI shows method-dependent behavior: comp_sim and strat_all favor small *k*, while strat_reduced exhibits an optimum at much larger *k*. The larger DBI variability for the β-heptapeptide is also consistent with its more heterogeneous conformational ensemble, where small differences in initial seeds can shift how sparsely populated or transitional conformations are partitioned. Accordingly, we use the conservative consensus range for subsequent β-heptapeptide analysis. The HP35 trajectory exhibits stronger agreement overall: both DBI and CHI consistently point to 5 to 7 clusters across seeding strategies, mirroring the values reported in our original *k*-Means NANI study.

**Table 1.** Preferred numbers of clusters for β-heptapeptide and HP35 based on the global DBI minimum and CHI maximum from the NANI screening.

| Seeding strategy | β-heptapeptide | HP35 |
| --- | --- | --- |
| **minimum DBI / 2<sup>nd</sup> minimum DBI** |  |  |
| *comp_sim* | 6/5 | 7/6 |
| *strat_all* | 5/25 | 7/6 |
| *strat_reduced* | 30/29 | 7/6 |
| **maximum CHI / 2<sup>nd</sup> maximum CHI** |  |  |
| *comp_sim* | 5/6 | 5/6 |
| *strat_all* | 5/6 | 5/6 |
| *strat_reduced* | 5/6 | 5/6 |

On the other hand, the local-curvature tests (Table S1) frequently wander to much larger values, especially for the β-heptapeptide, confirming earlier observations that second-derivative metric can also over-react to minor fluctuations in the curves for these systems. Because the global metrics are both more conservative and more consistent across seeding modes, we retained them as the principal guide and fixed *k* = 6 for β-heptapeptide and *k* = 7 for HP35, a choice also supported by our previous analysis of this system.^7^ The following section explores how these NANI clusters partition the conformational landscape of β-heptapeptide and HP35 and compare their properties as we vary the initialization schemes.

To verify that strat_all and strat_reduced recover the same physical states as comp_sim at the optimal cluster count, we first analyzed the β-heptapeptide at *k* = 6 (Table 2). Overall, the three strategies yield closely related partitions, with the main differences reflecting how peripheral frames are distributed among clusters. Relative to comp_sim, the stratified seeders shift population away from the most populated cluster and reduce its MSD, consistent with a tighter core. At the same time, the lower-population clusters increase in size and exhibit higher MSD values, indicating that the additional frames lie farther from their cluster centers. Although the medoids are not identical, the same key conformational states are recovered for this system. Fig. 3 shows representative structures for the six β-heptapeptide clusters obtained with NANI seeded using strat_reduced.

**Table 2.** Cluster populations and MSD for β-heptapeptide with *k* = 6 clusters obtained with the thre NANI modes. Clusters are ordered in increasing population.

| comp_sim |  | strat_all |  | strat_reduced |  |
| --- | --- | --- | --- | --- | --- |
| Population | MSD | Population | MSD | Population | MSD |
| 582 | 7.33 | 680 | 7.44 | 663 | 7.33 |
| 645 | 4.15 | 738 | 4.85 | 741 | 4.86 |
| 693 | 7.76 | 946 | 10.58 | 1007 | 10.62 |
| 1012 | 10.83 | 951 | 10.46 | 1067 | 10.39 |
| 1225 | 9.68 | 1235 | 9.13 | 1210 | 9.27 |
| 1844 | 10.35 | 1451 | 9.69 | 1313 | 9.51 |

**Figure 3.**
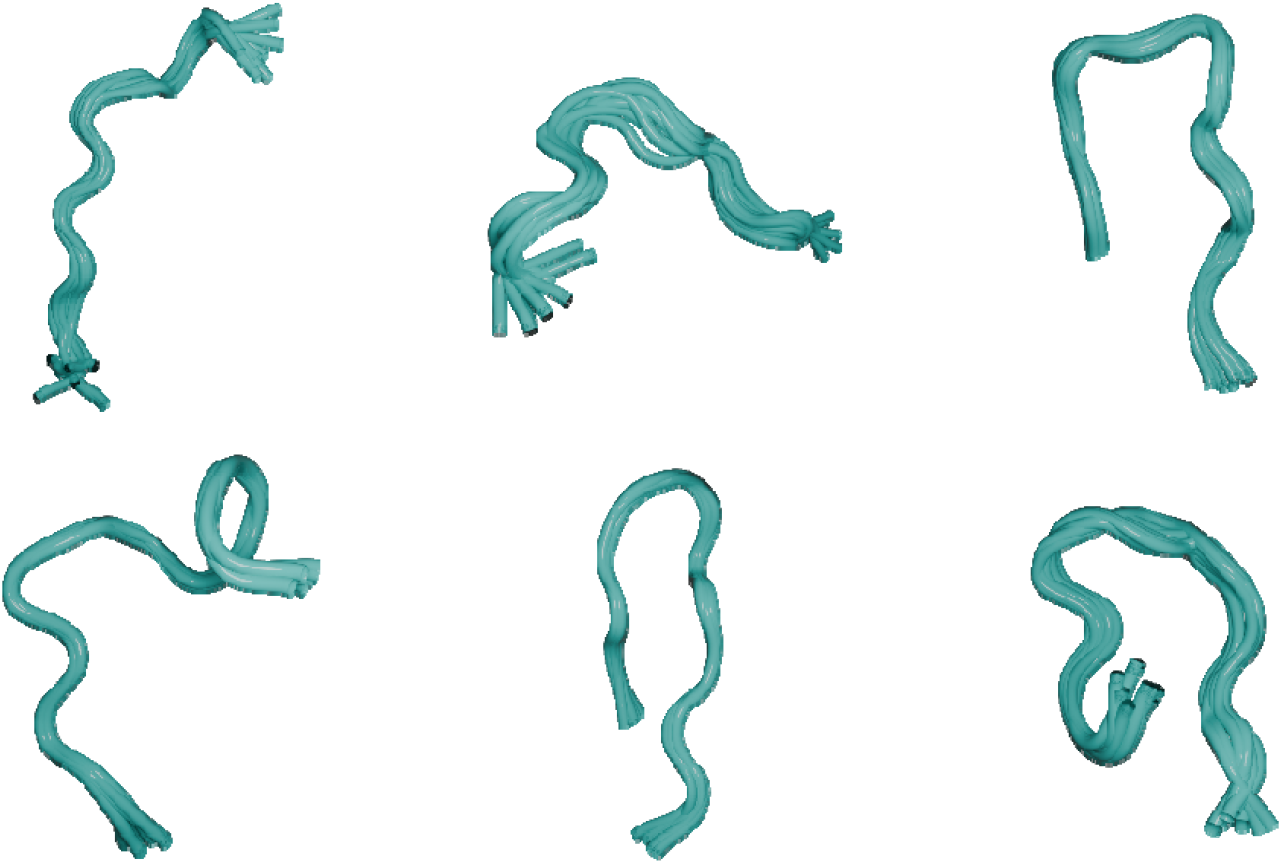
Representative structures for the six β-heptapeptide clusters obtained with NANI seeded using strat_reduced. Representative structures are shown as the medoid plus the ten closest frames to the medoid (11 structures per cluster).

We also examined the cluster populations and MSDs for HP35 at *k* = 7. As Table 3 shows, the seven clusters are virtually unchanged by the choice of seed: each cluster’s population differs by fewer than 40 structures out of ∼150k frames (< 0.03 %), and the MSDs vary by at most 0.15. Fig. 4 shows the best representative frames from each cluster produced by strat_reduced, illustrating the clear separation and compactness of the resulting clusters. Upon inspection, we found that the same representative structures were selected by comp_sim and strat_all as well. In other words, all three initialization methods produced identical medoid frames at *k* = 7. Together, these results demonstrate that the new stratification methods carve out the same conformational states, with the same structural heterogeneity as the original protocol, but at a fraction of the computational cost.

**Table 3.** Cluster populations and MSD for HP35 with *k* = 7 clusters obtained with the three NANI modes. Clusters are ordered in increasing population.

| comp_sim |  | strat_all |  | strat_reduced |  |
| --- | --- | --- | --- | --- | --- |
| Population | MSD | Population | MSD | Population | MSD |
| 5577 | 80.44 | 5569 | 80.33 | 5577 | 80.44 |
| 8123 | 91.31 | 8085 | 91.37 | 8121 | 91.31 |
| 11929 | 76.01 | 11950 | 76.01 | 11927 | 76.02 |
| 12096 | 46.55 | 12108 | 46.70 | 12100 | 46.55 |
| 13274 | 72.29 | 13260 | 72.29 | 13274 | 72.29 |
| 32815 | 14.10 | 32824 | 14.11 | 32815 | 14.10 |
| 67291 | 9.27 | 67309 | 9.28 | 67291 | 9.27 |

**Figure 4.**
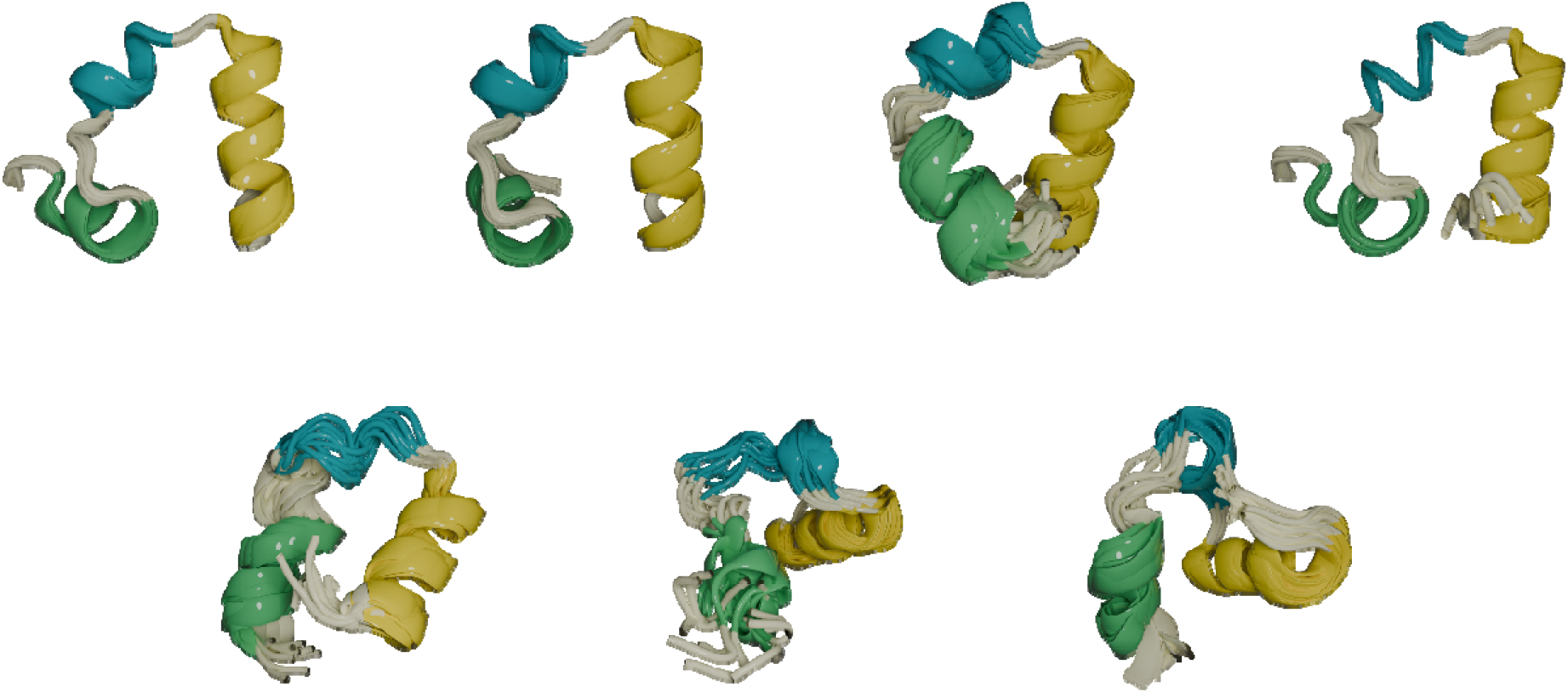
Representative structures for the seven HP35 clusters obtained with NANI seeded using strat_reduced. Representative structures are shown as the medoid plus the ten closest frames to the medoid. Helices 1, 2, and 3 are colored green, cyan, and yellow, respectively.

Because the representative (medoid) frame indices are identical across methods for HP35, cluster correspondence is direct, whereas for the β-heptapeptide representative indices and population ordering differ and clusters are not assumed to align by index.

### 4.2. Runtime and scaling performance

We next quantified the speed advantage of the new strategies by benchmarking a full *k-*means pass at *k* = 60. The results in Table 4 show that the stratified seeders already save approximately 30% of the runtime on a small β-heptapeptide trajectory and scale far better on the larger HP35 dataset, where the time spent on the complete cycle dramatically decreased from almost 30 min to ∼160 s, an order-of-magnitude improvement. In practical terms, the stratified seeders deliver clusters of similar quality at about one-tenth the cost on a 150k frame trajectory. The payoff becomes more dramatic at larger scales: *k*-means clustering of the full 1.5 million HP35 frames at *k* = 60 using comp_sim took ∼30 hours whereas strat_all and strat_reduced now complete the same task in ∼40 min, almost a 45-fold speedup.

**Table 4.**
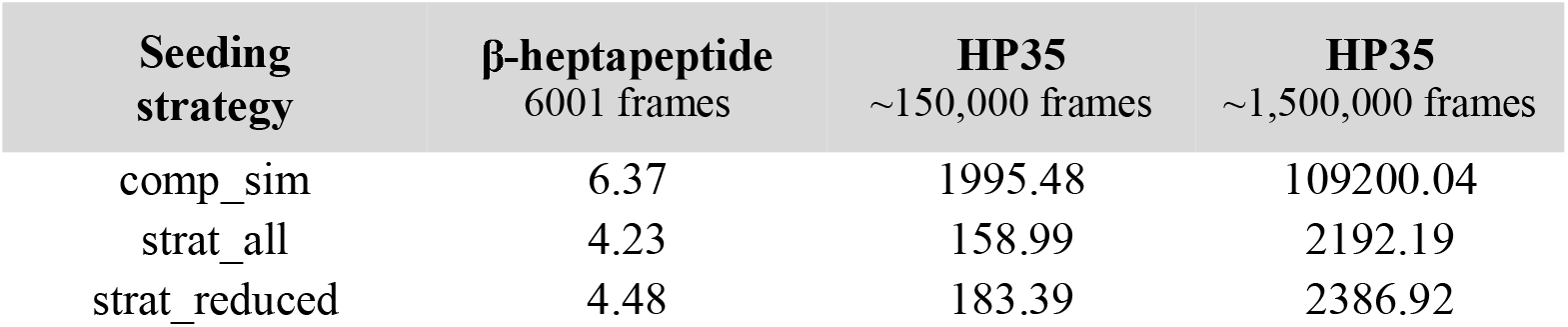
Runtime (s) to complete a single *k-*means pass (*k* = 60) with each NANI seeding strategy.

### 4.3. Effect of frame count and representation size

To isolate effect of representation size to the effect of the number of frames, we performed timing scans over increasing frame counts for both benchmark systems and fit a linear model, where the slope provides an empirical per-frame cost (s/frame). For these scans we used *k* = 6 for the β-heptapeptide system and *k* = 7 for HP35, consistent with the cluster counts used in the cluster analysis. The β-heptapeptide representation contained 50 atoms (150 features), while HP35 contained 101 atoms (303 features). Fig. 5 shows elapsed time as a function of frame count, and Table 5 reports the fitted slopes and goodness-of-fit values.

**Table 5.** Linear regression summary for the scaling analysis. For each strategy, the fitted slope and the coefficient of determination are reported.

| System | Seeding strategy | Slope (s/frame) | $R^2$ |
| --- | --- | --- | --- |
| β-heptapeptide | comp_sim | | 0.95 |
|  | strat_all |  | 0.90 |
|  | strat_reduced |  | 0.88 |
| HP35 | comp_sim |  | 0.95 |
|  | strat_all |  | 0.67 |
|  | strat_reduced |  | 0.80 |

**Figure 5.**
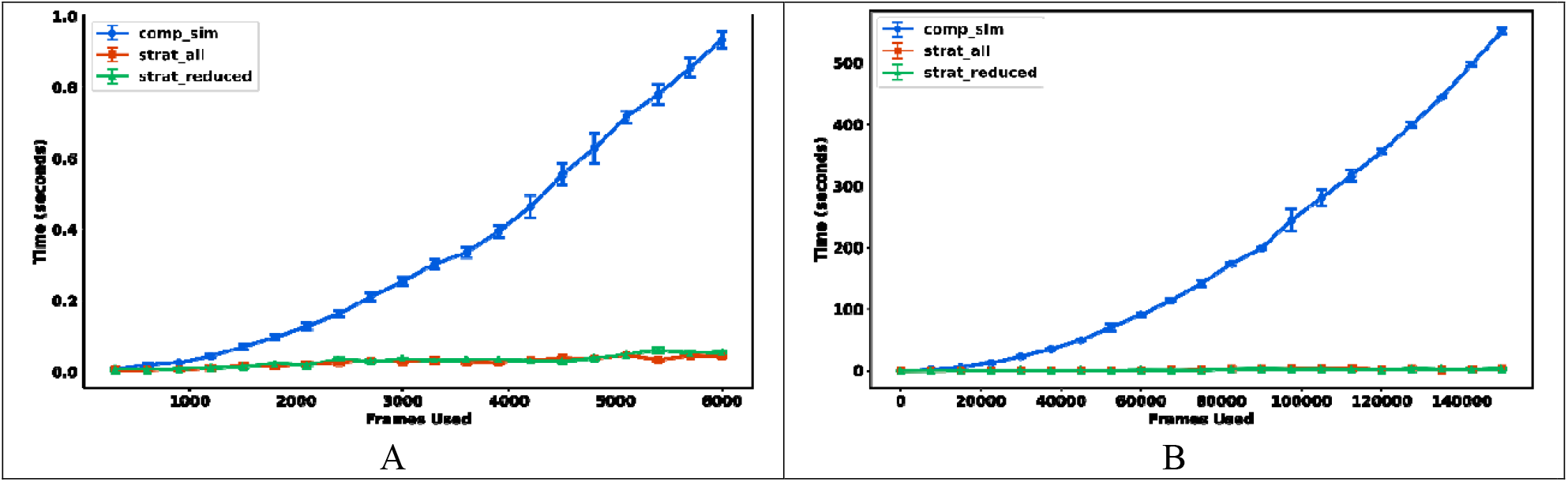
Runtime as a function of the number of frames for (A) β-heptapeptide (*k* = 6; 50 atoms) and (B) HP35 (*k* = 7; 101 atoms). Points show mean elapsed time over three replicate scans; error bars show the standard deviation.

Across both systems, strat_all and strat_reduced exhibited near-linear scaling with frame count and substantially smaller slopes than comp_sim, indicating that runtime grows slowly with dataset size. The fitted slopes also quantify sensitivity to representation size. Increasing the clustering representation from 50 atoms (β-heptapeptide) and 101 atoms (HP35) increases the per-frame slope by ∼3.3x for strat all and ∼2.7x for strat_reduced, whereas comp_sim increased by ∼24x. This shows that the stratified initializations are substantially less sensitive to system size than comp_sim, consistent with their one-pass design.

### 4.4. Integration in the HELM workflow

In the Hierarchical Extended Linkage Method (HELM)^22^ pipeline, NANI provides a deterministic *k-* means pre-clustering that groups frames into small representative clusters before the hierarchical merging. In our previous runs, this pre□clustering, performed with the comp_sim variant of NANI, dominated the computational cost. On the 151k HP35 frames, the NANI part consumed ∼33□min, accounting for 98□% of HELM’s total runtime, creating a clear bottleneck. Replacing comp_sim with either strat_all or strat_reduced eliminates this challenge. A complete HELM run now finishes in ∼3 min, a full order-of-magnitude speed□up (Table S2).

We saw that the significant speed-up does not compromise on clustering quality. While each seeding strategy produced a different set of initial *k*-means cluster labels, running HELM on these different inputs yielded highly similar results. As we show on Fig. 6, the CHI and DBI at each merging step in the HP35 system follow the same trends for all three initialization methods. Fig. 7 presents the overlaps of the best representative frames when *k* = 6. This improvement now makes fully deterministic clustering feasible for the multi-million frame trajectories typical of modern large-scale MD simulations.

**Figure 6.**
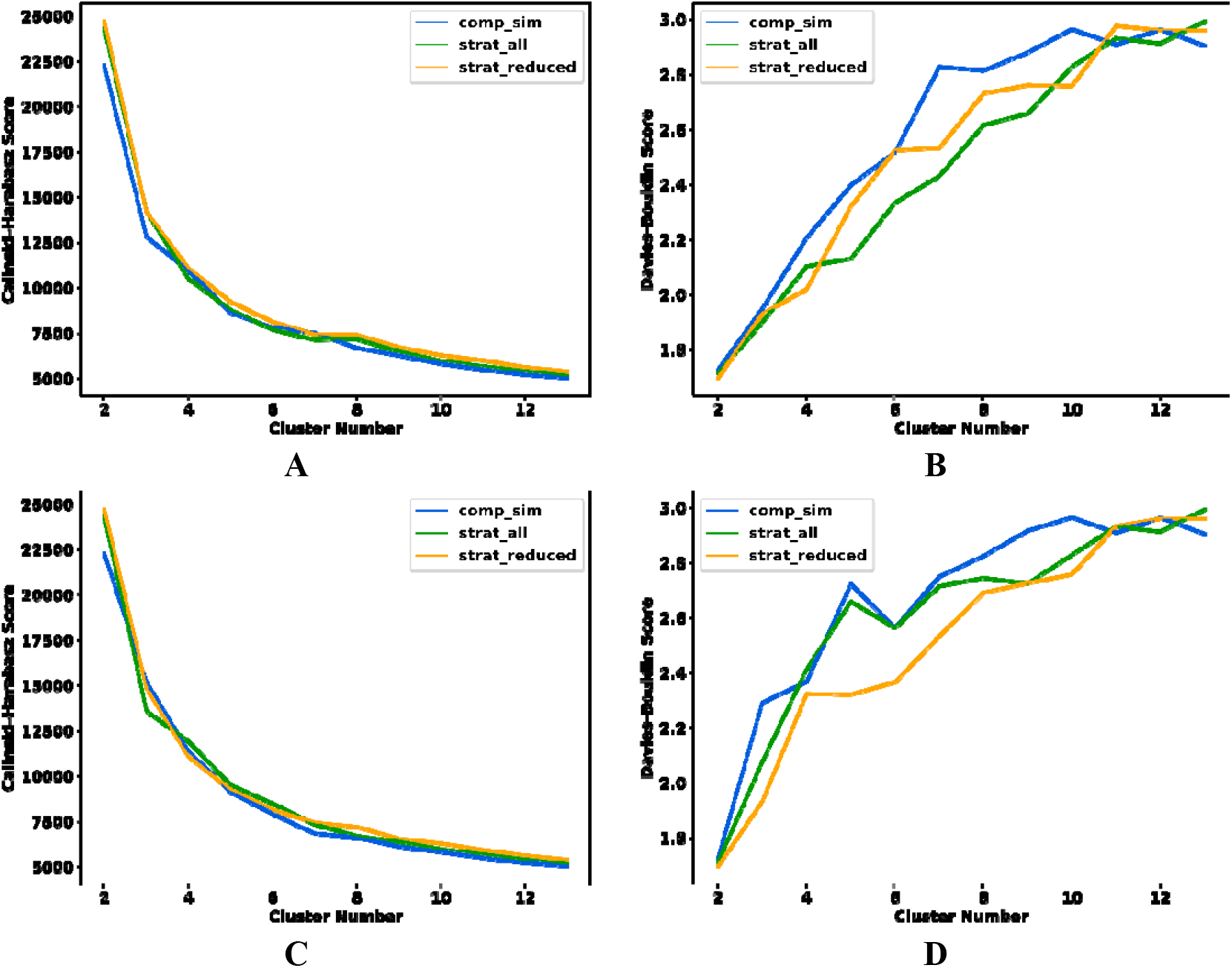
Change in Calinski-Harabasz (A, C) and Davies-Bouldin (B, D) indices for the HP35 simulation after trimming the initial NANI clusters (retaining only clusters with MSD < 10) using the (A, B) inter- and the (C, D) intra-linkage.

**Figure 7.**
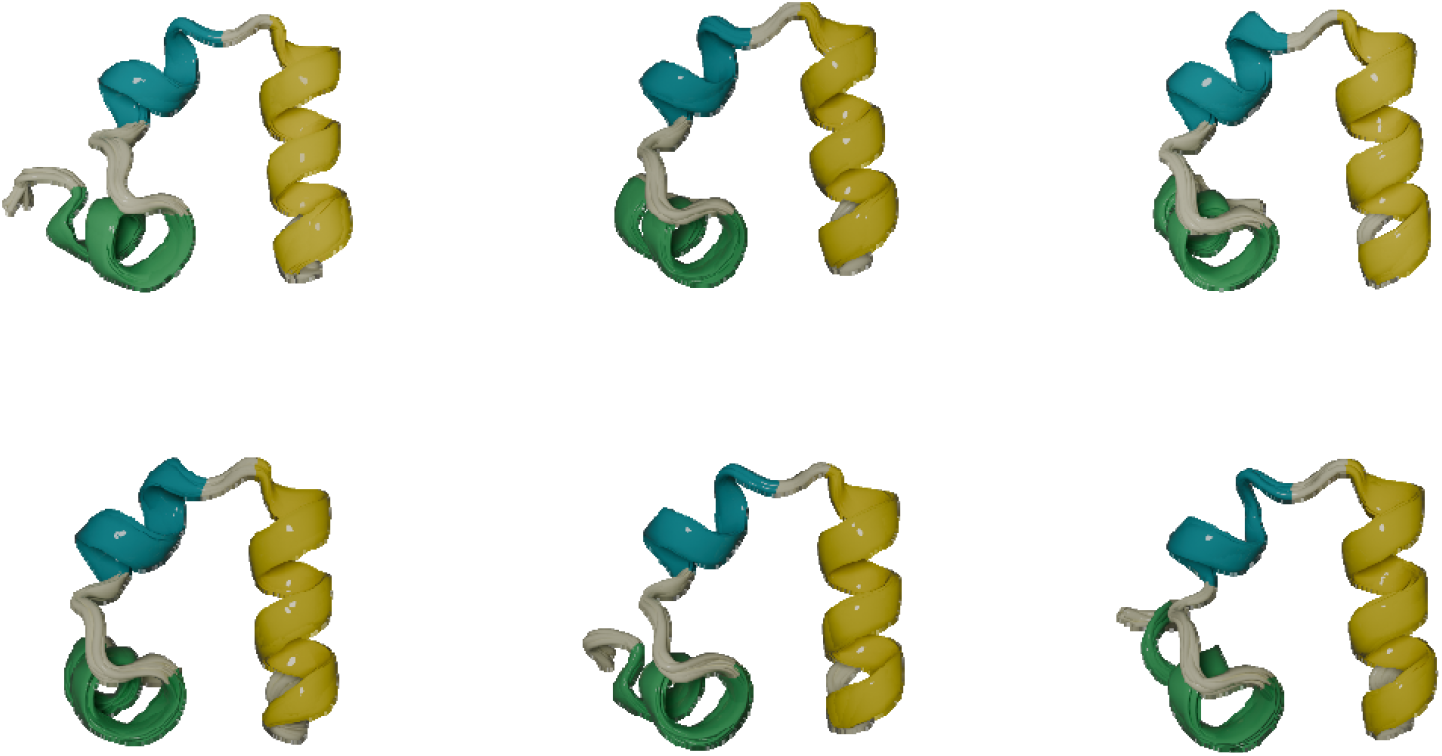
Representative structures for the six final HP35 clusters obtained with HELM (intra-linkage) starting from 60 NANI pre-clusters seeded with strat_reduced. HELM was run with trimming (retaining only clusters with MSD < 10)

## 5. CONCLUSION

We have presented two deterministic, stratified *k*-means initialization strategies: strat_all and strat_reduced, that improve on the efficiency of *k*-means NANI clustering for MD simulations. These new variants address a key limitation of the previous NANI implementations: the high computational cost of iterative seed selection on large datasets. By replacing that step with a one-pass stratified sampling based on complementary similarity, we retain the core strengths of the method: determinism, robustness, and physical interpretability, while dramatically reducing runtime. Our benchmarks across systems demonstrate that these improvements do not sacrifice clustering quality, as evidenced by the consistent CHI and DBI scores as well as cluster populations that are in agreement with those from prior NANI runs.

For routine use, we recommend strat_all or strat_reduced as the default *k*-means NANI initialization because they preserve clustering quality while reducing initialization cost by orders of magnitude at large-scale MD. The original comp_sim strategy can still be beneficial when users want to restrict seeds to the highest-density subset of the frames or when working with small datasets where the additional cost is negligible and the iterative diversity selection may provide slightly different partitions. In practice, we recommend strat_reduced when the goal is to emphasize dense-state coverage by restricting seeds to high-cMSD frames, and strat_all when broader coverage of the full ensemble is desired.

Importantly, we showed how these new strategies substantially improve the efficiency of HELM. The original HELM pipeline leveraged NANI pre-clustering to group frames into small clusters before the hierarchical merging. However, as noted in our prior work, the NANI step had become the computational bottleneck, particularly on large-scale trajectories. By replacing comp_sim with strat_all or strat_reduced, we removed this bottleneck entirely. Clustering runtimes fell by more than an order of magnitude while producing hierarchies that preserved the biophysically interpretable clusters.

Again, the success of these new stratified NANI methods lies not only in their significant acceleration of the seed selection process in *k*-means clustering but also in how they enable scalable downstream hierarchical workflows. By replacing greedy initialization with deterministic stratification, our methods remove one of the key barriers to utilizing hybrid protocols like HELM at large scale. As such, strat_all and strat_reduced extend NANI’s impact beyond standalone use, positioning it as an important tool for advancing routine, scalable, and reproducible analysis of complex conformational ensembles in molecular simulations.

## SUPPORTING INFORMATION

Second-derivative cluster-count analysis for β-heptapeptide and HP35; representative cluster structures for NANI and HELM workflows; DBI sensitivity to the stratification percentage parameter *p*; RMSD-to-first-frame traces and distributions; and additional runtime benchmarks for NANI and HELM processing (PDF).

## ACKNOWLEDGEMENTS

RAMQ, LC, and JBWS thank support from the National Institute of General Medical Sciences of the National Institutes of Health under award number R35GM150620.

## AUTHOR CONTRIBUTIONS

JBWS: Data curation; Formal analysis; Investigation; Software; Validation; Visualization; Writing. LC: Data curation; Formal analysis; Investigation; Software; Validation; Visualization; Writing.

RAMQ: Formal analysis; Methodology; Conceptualization; Investigation; Software; Writing; Funding acquisition; Supervision; Resources.

## DATA AND SOFTWARE AVAILABILITY

The NANI code can be found here: https://github.com/mqcomplab/MDANCE.

## TOC Graphic

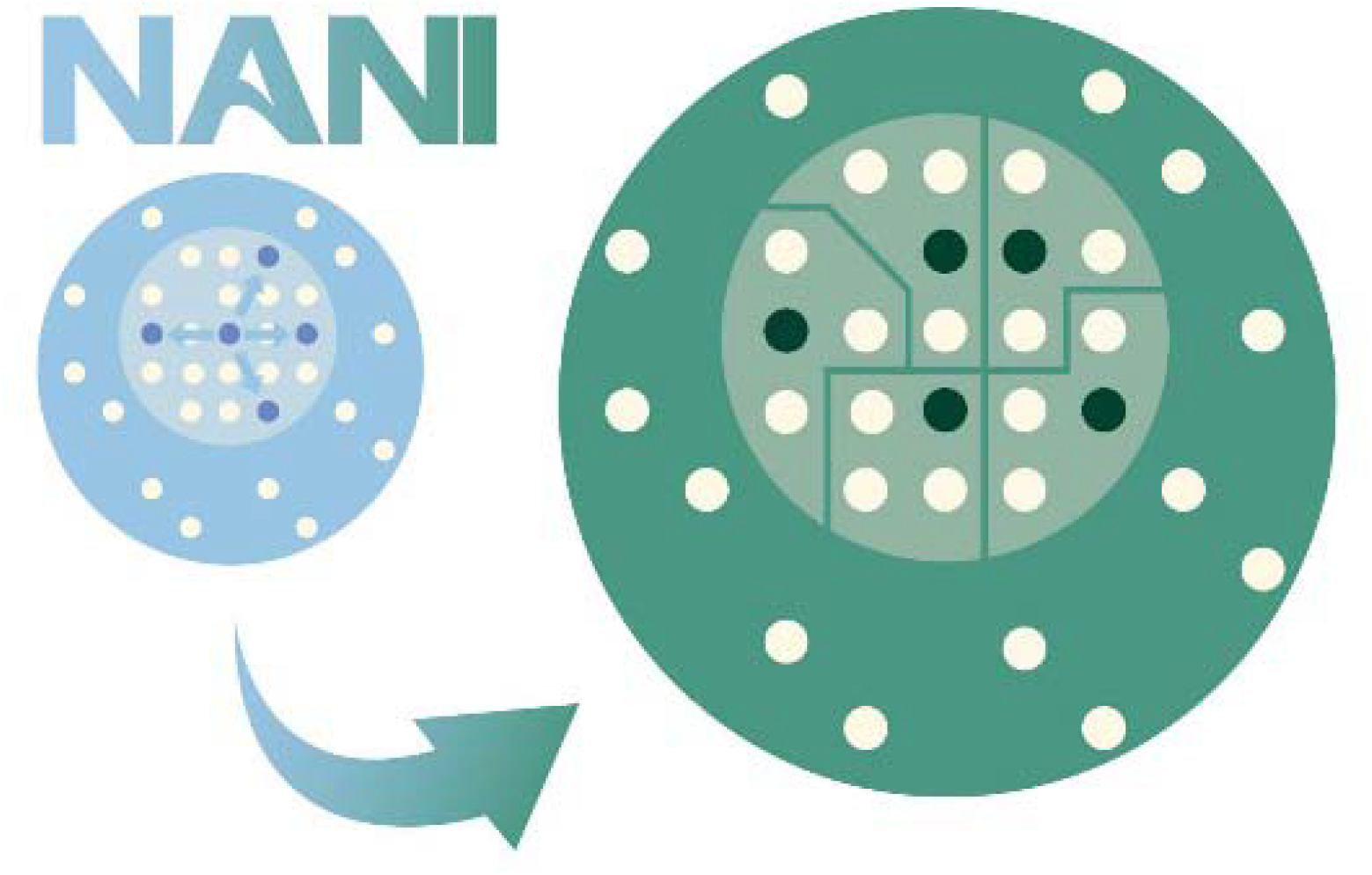

