## Supplementary Information for "Scaling *k*-Means for Multi-Million Frames: A Stratified NANI Approach for Large-Scale MD Simulations"

**Table S1:** Preferred numbers of clusters for β-heptapeptide and HP35 based on the local second-derivative extrema metrics after the NANI screening. Blank cells indicate that no local minimum met the criteria.

| **Seeding strategy** | **β-heptapeptide** | **HP35** |
| --- | --- | --- |
| **Maximum 2^nd^ Derivative of DBI** | | |
| comp_sim | 8 | 16 |
| strat_all | 25 | 11 |
| strat_reduced | 9 | 11 |
| **Minimum 2^nd^ Derivative of CHI** | | |
| comp_sim | - | - |
| strat_all | - | - |
| strat_reduced | 19 | - |

| 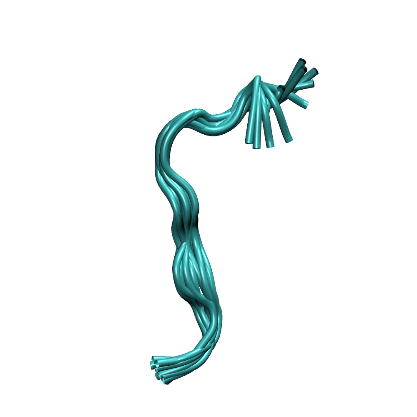**A1** | 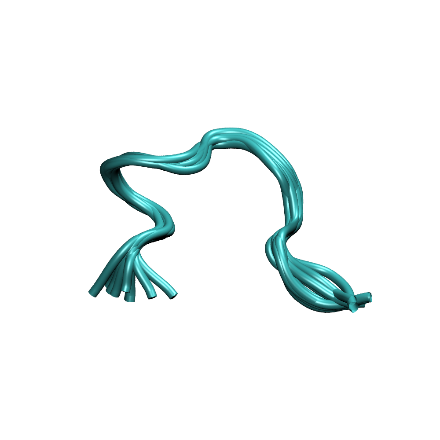**A2** | 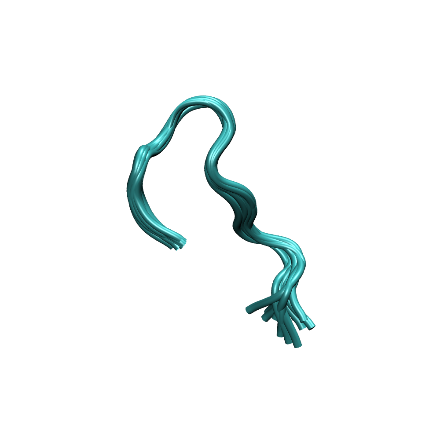**A3** |
| --- | --- | --- |
| 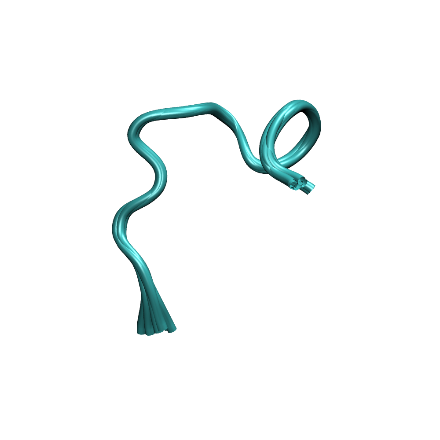**A4** | 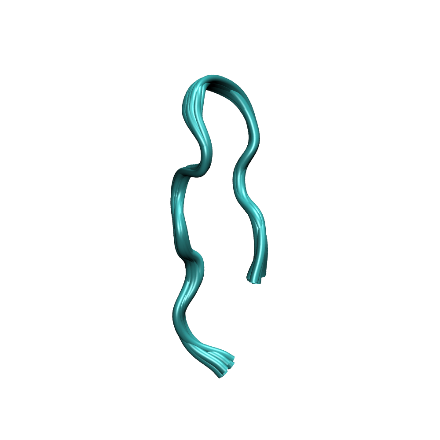**A5** | 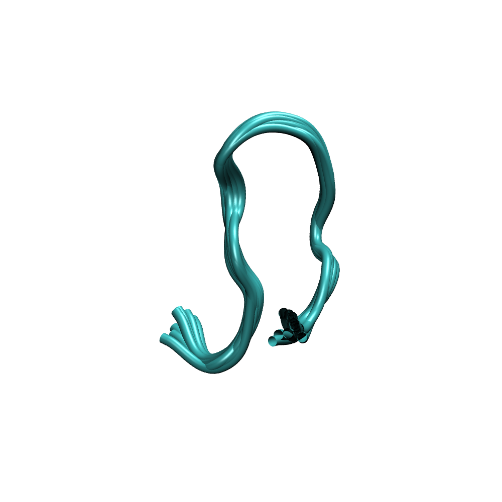**A6** |
| 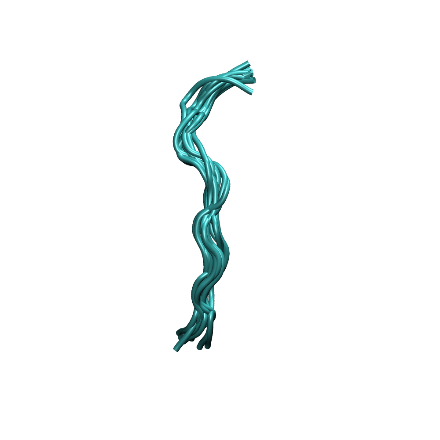**B1** | 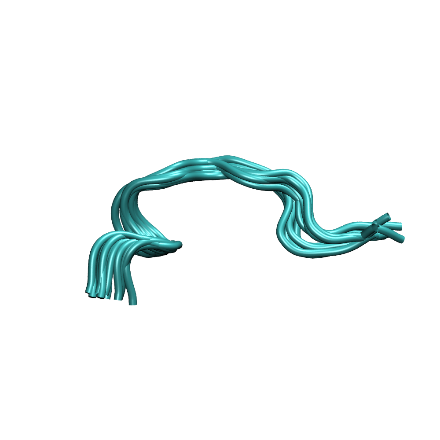**B2** | 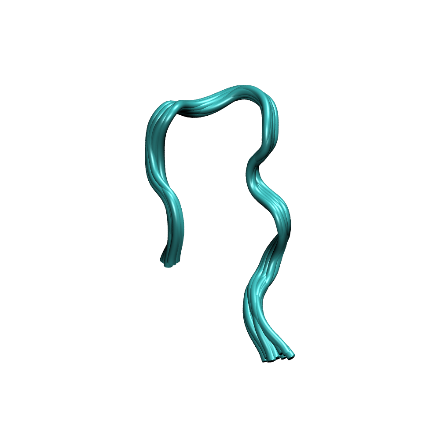**B3** |
| 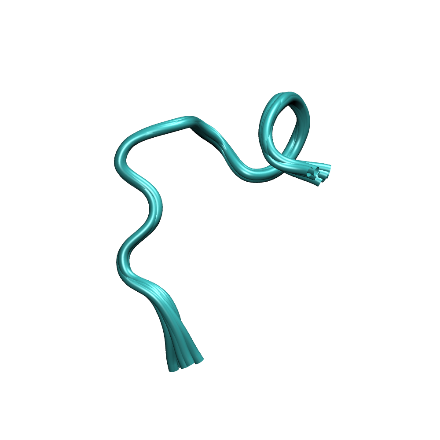**B4** | 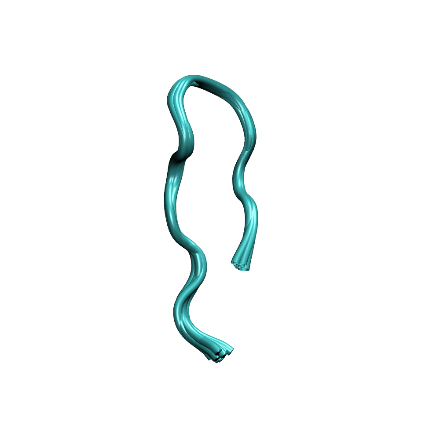**B5** | 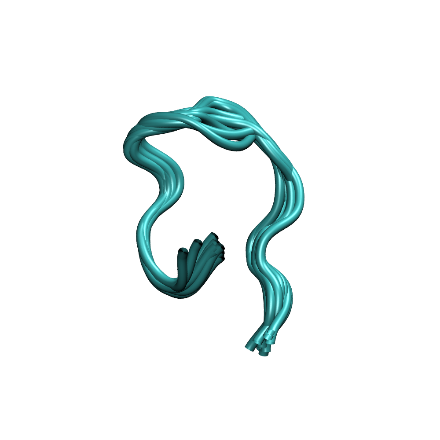**B6** |

**Figure S1.** Representative structures for the six β-heptapeptide clusters obtained with NANI seeded using comp_sim (A panels) and strat_all (B panels). Representative structures are shown as the medoid plus the ten closest frames to the medoid.

| 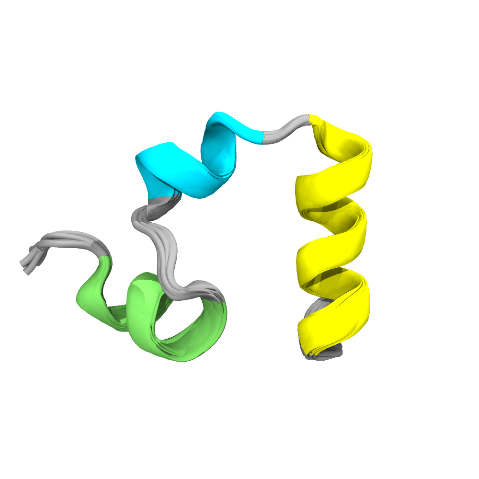**A1** | 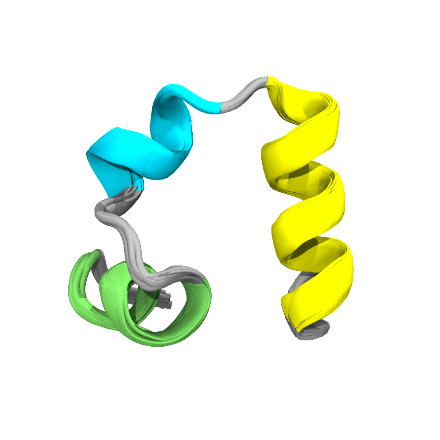**A2** | 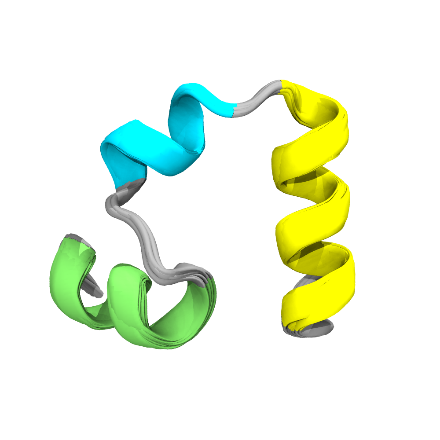**A3** |
| --- | --- | --- |
| 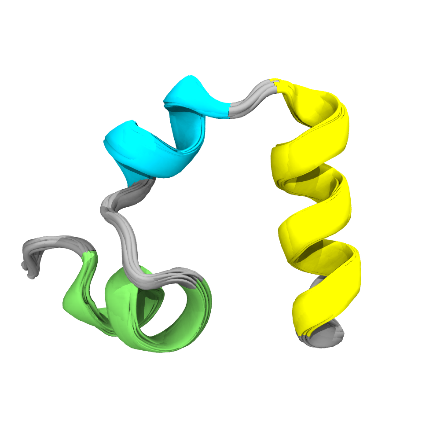**A4** | 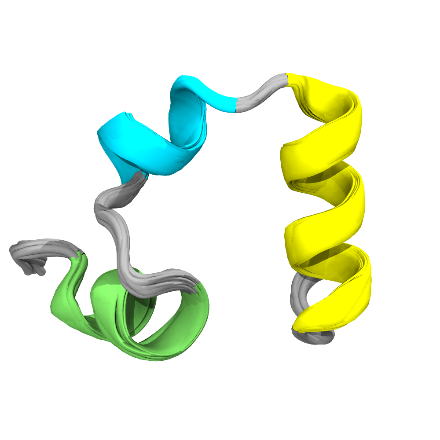**A5** | 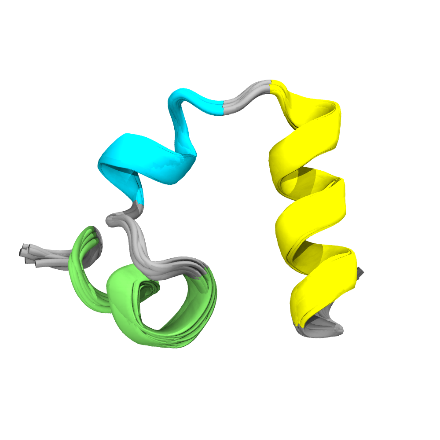**A6** |
| 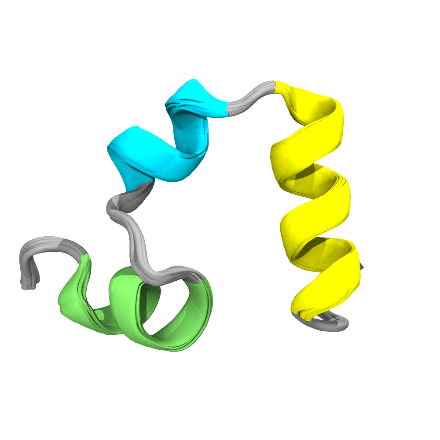**B1** | 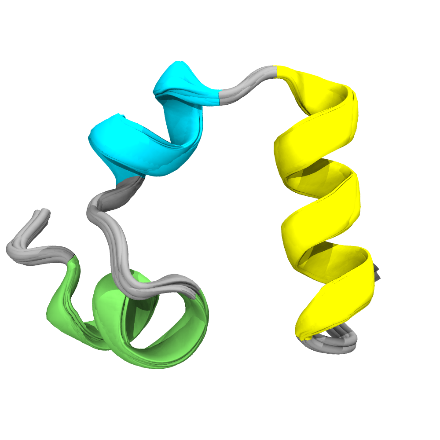**B2** | 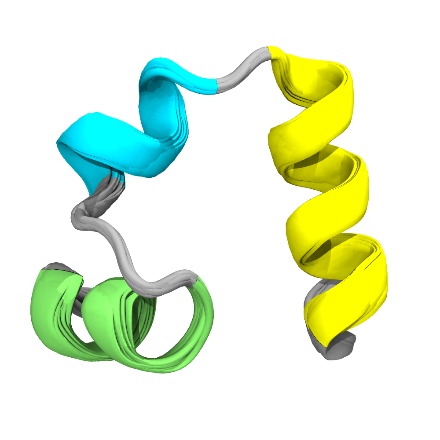**B3** |
| 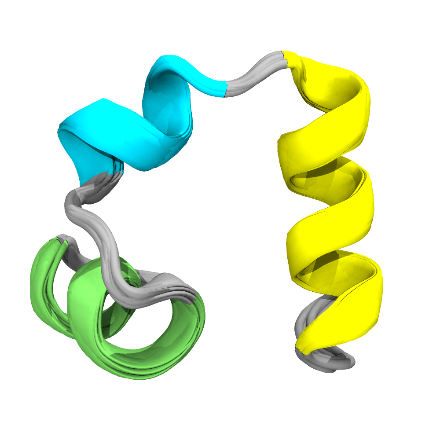**B4** | 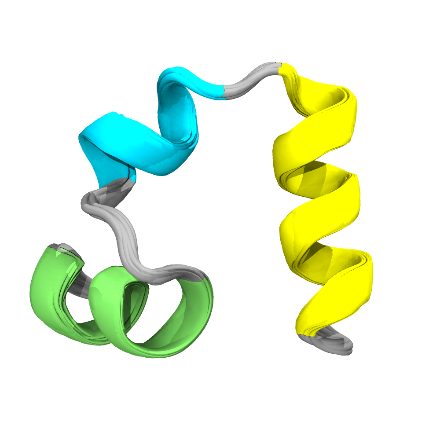**B5** | 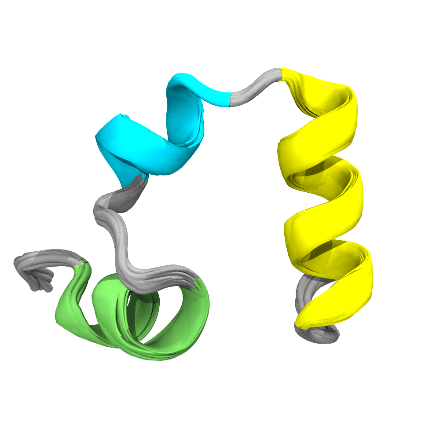**B6** |

**Figure S2.** Representative structures for the six final HP35 clusters obtained with HELM starting from 60 NANI pre-clusters seeded with comp_sim. Panels show inter-linkage (A) and intra-linkage (B). HELM was run with trimming (retain clusters with MSD < 10). Representative structures are shown as the medoid plus the ten closest frames to the medoid.

| 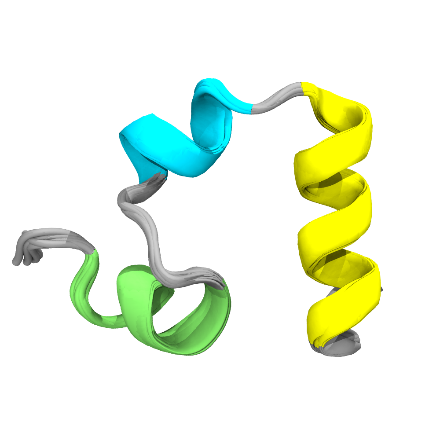**A1** | 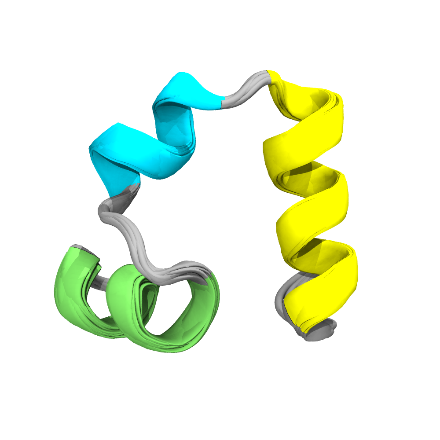**A2** | 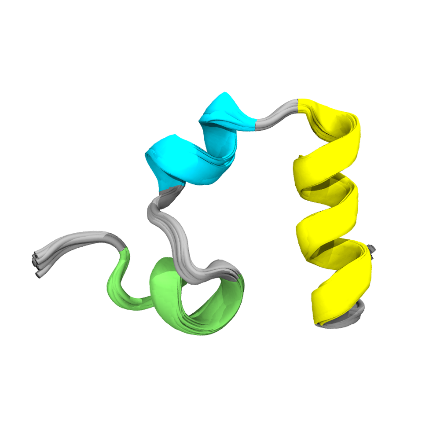**A3** |
| --- | --- | --- |
| 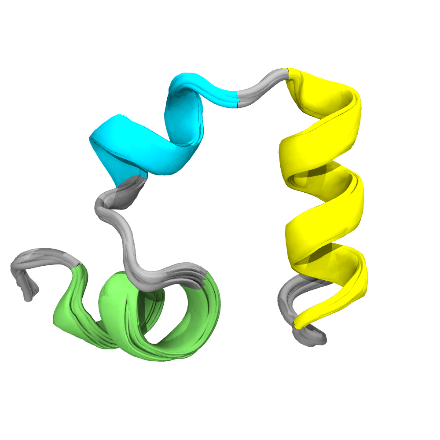**A4** | 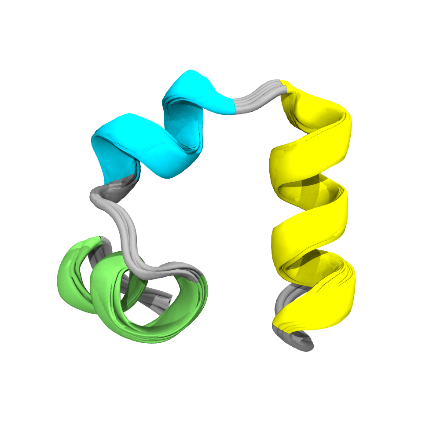**A5** | 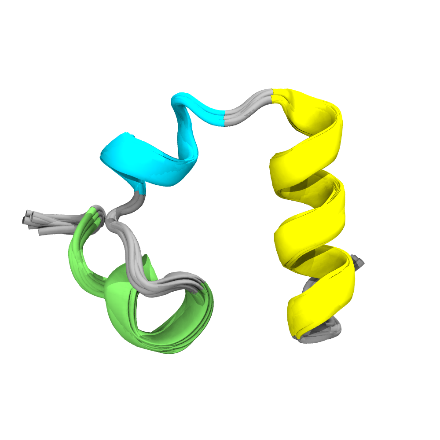**A6** |
| 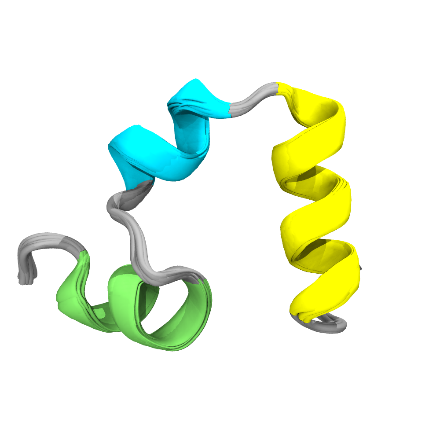**B1** | 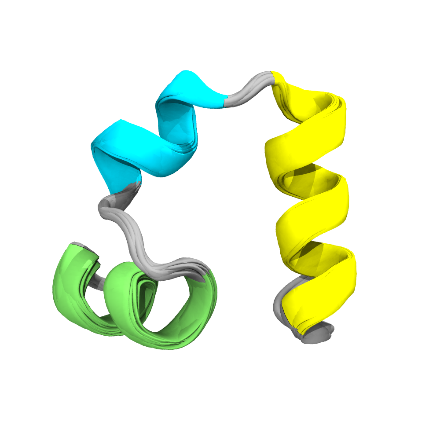**B2** | 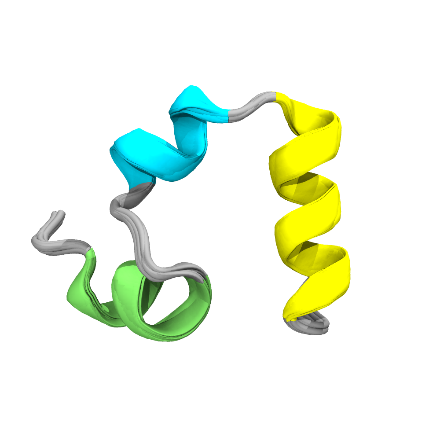**B3** |
| 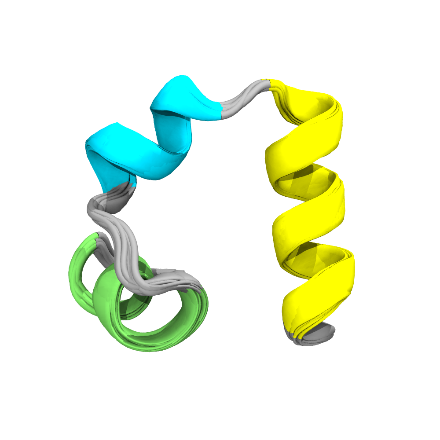**B4** | 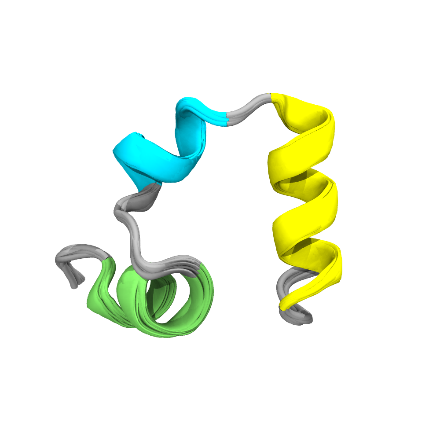**B5** | 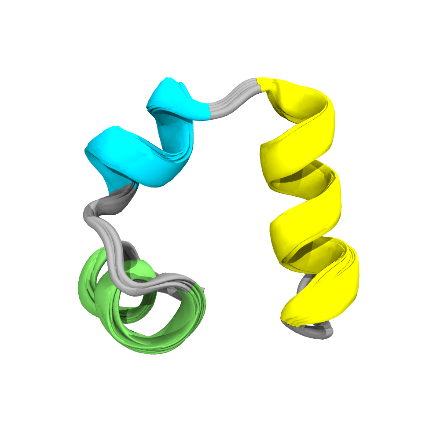**B6** |

**Figure S3.** Representative structures for the six final HP35 clusters obtained with HELM starting from 60 NANI pre-clusters seeded with strat_all. Panels show inter-linkage (A) and intra-linkage (B). HELM was run with trimming (retain clusters with MSD < 10). Representative structures are shown as the medoid plus the ten closest frames to the medoid.

| 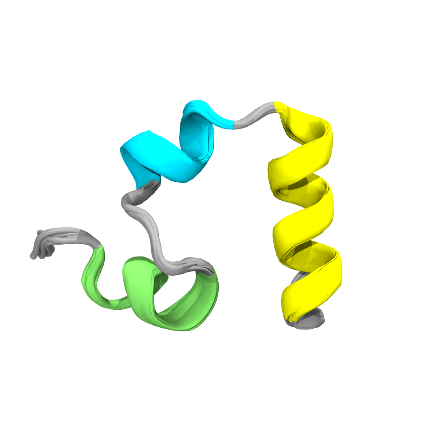**A1** | 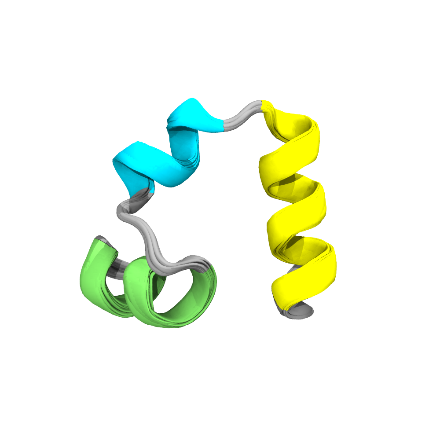**A2** | 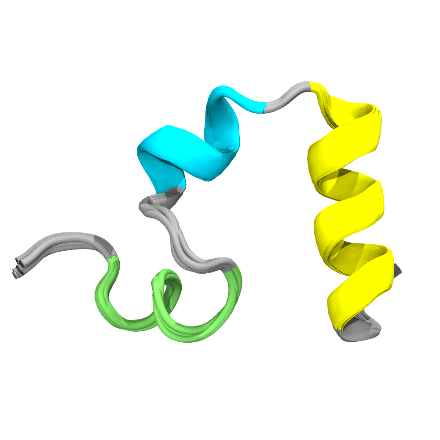**A3** |
| --- | --- | --- |
| 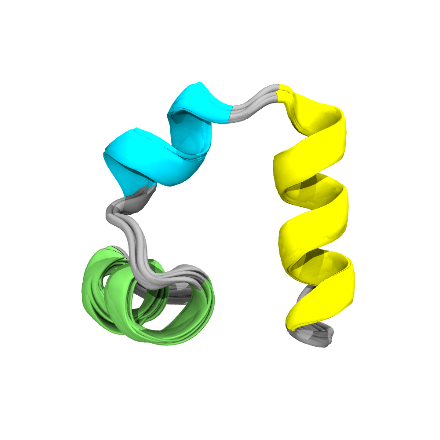**A4** | 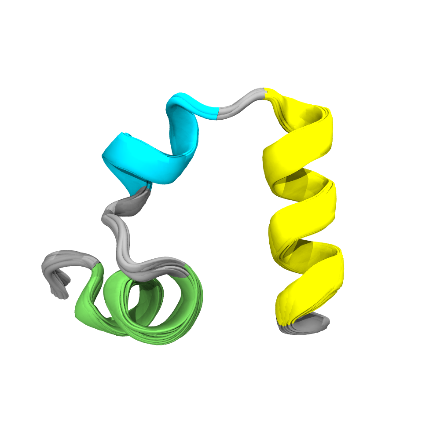**A5** | 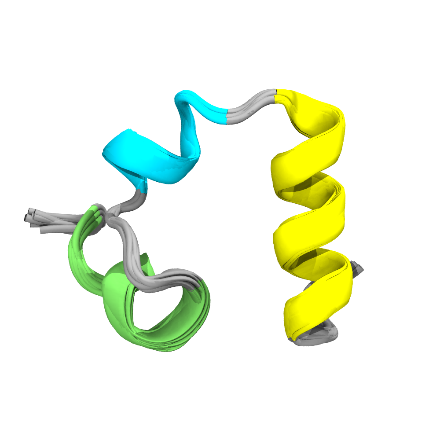**A6** |

**Figure S4.** Representative structures for the six final HP35 clusters obtained with HELM starting from 60 NANI pre-clusters seeded with strat_reduced using the inter-linkage merge scheme. HELM was run with trimming (retain clusters with MSD < 10). Representative structures are shown as the medoid plus the ten closest frames to the medoid.

**Table S2:** Runtime (s) to complete NANI and HELM processing of HP35 (sieve = 10) with trimming (retaining clusters with MSD < 10). The NANI time includes the *k*-means initialization and clustering, while the HELM time includes the hierarchical merging.

| **method** | **Linkage** | **NANI Time (s)** | **HELM Time (s)** | **Total (s)** |
| --- | --- | --- | --- | --- |
| comp_sim | inter | 1995.48 | 8.41 | 2003.89 |
|  | intra |  | 11.06 | 2006.54 |
| strat_all | inter | 158.99 | 6.05 | 165.04 |
|  | intra |  | 11.29 | 170.28 |
| strat_reduced | inter | 183.39 | 9.42 | 192.81 |
|  | intra |  | 10.78 | 194.17 |

| 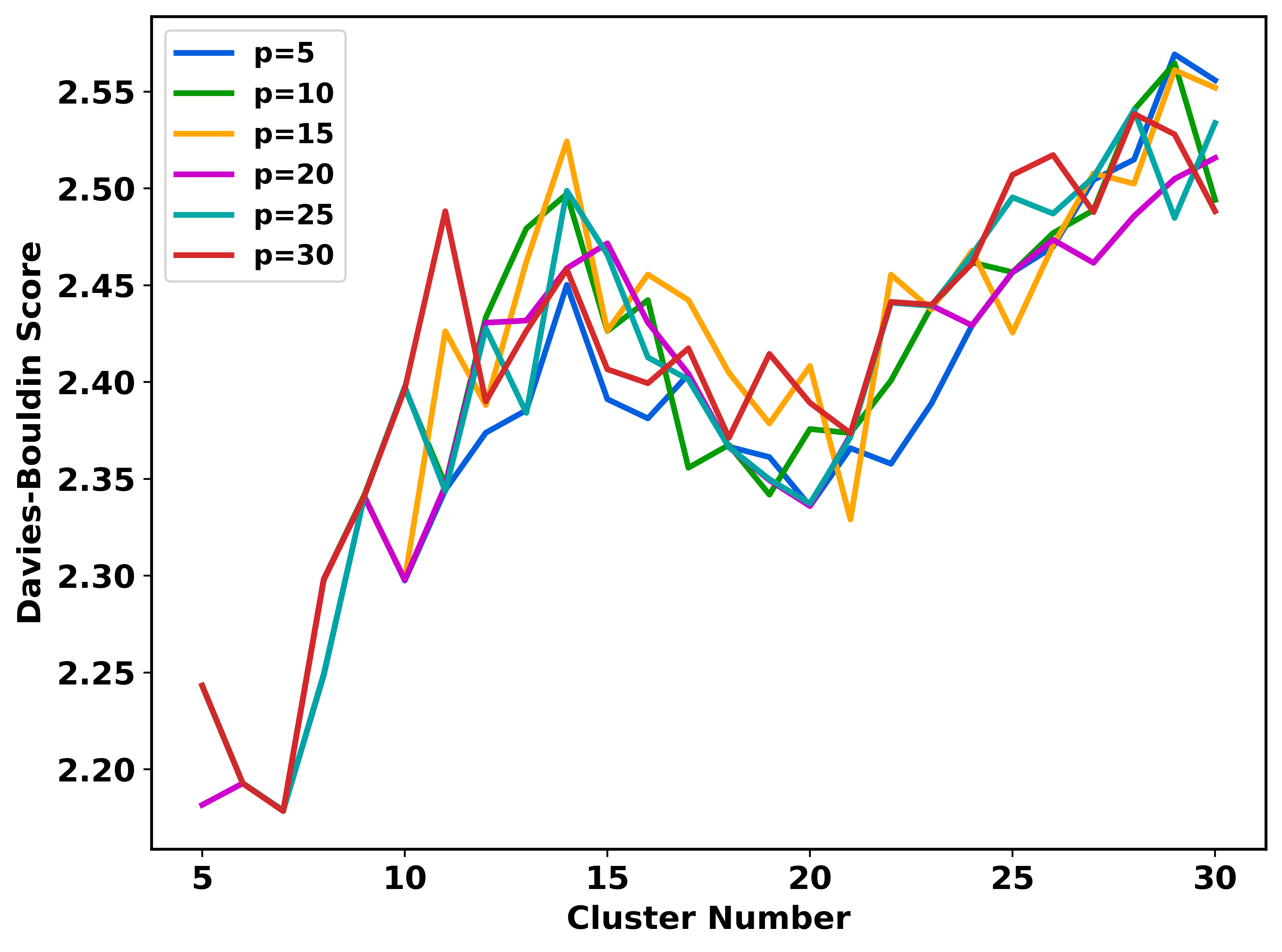  A | 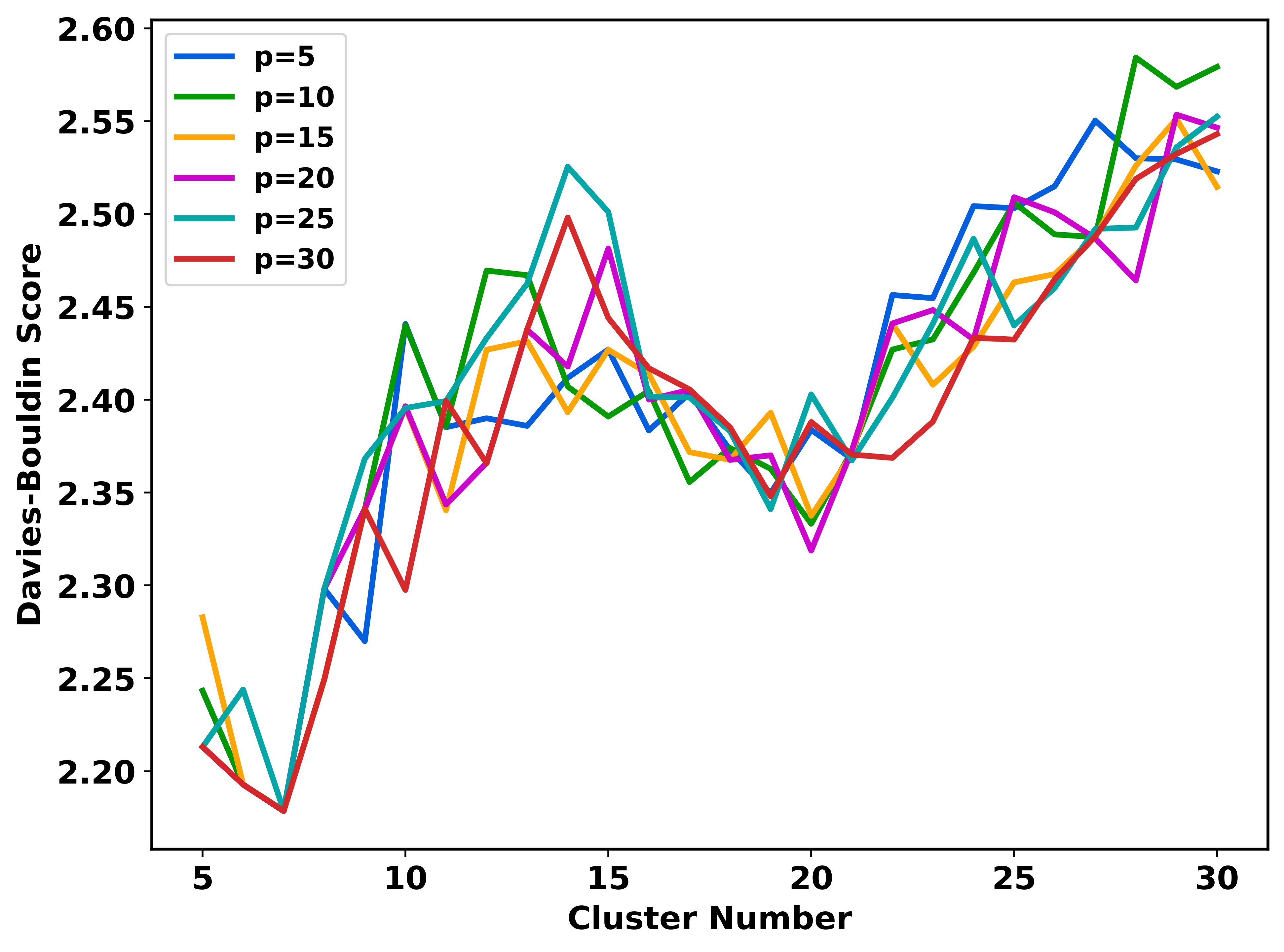  B |
| --- | --- |

**Figure S5.** Sensitivity of (A) strat_all and (B) strat_reduced initialization to the percentage parameter *p* on the HP35 system. DBI values across clustering levels for *p* = 5% to 30%. DBI trends remain consistent across the tested range, indicating limited dependence of clustering quality on *p* for the system evaluated.

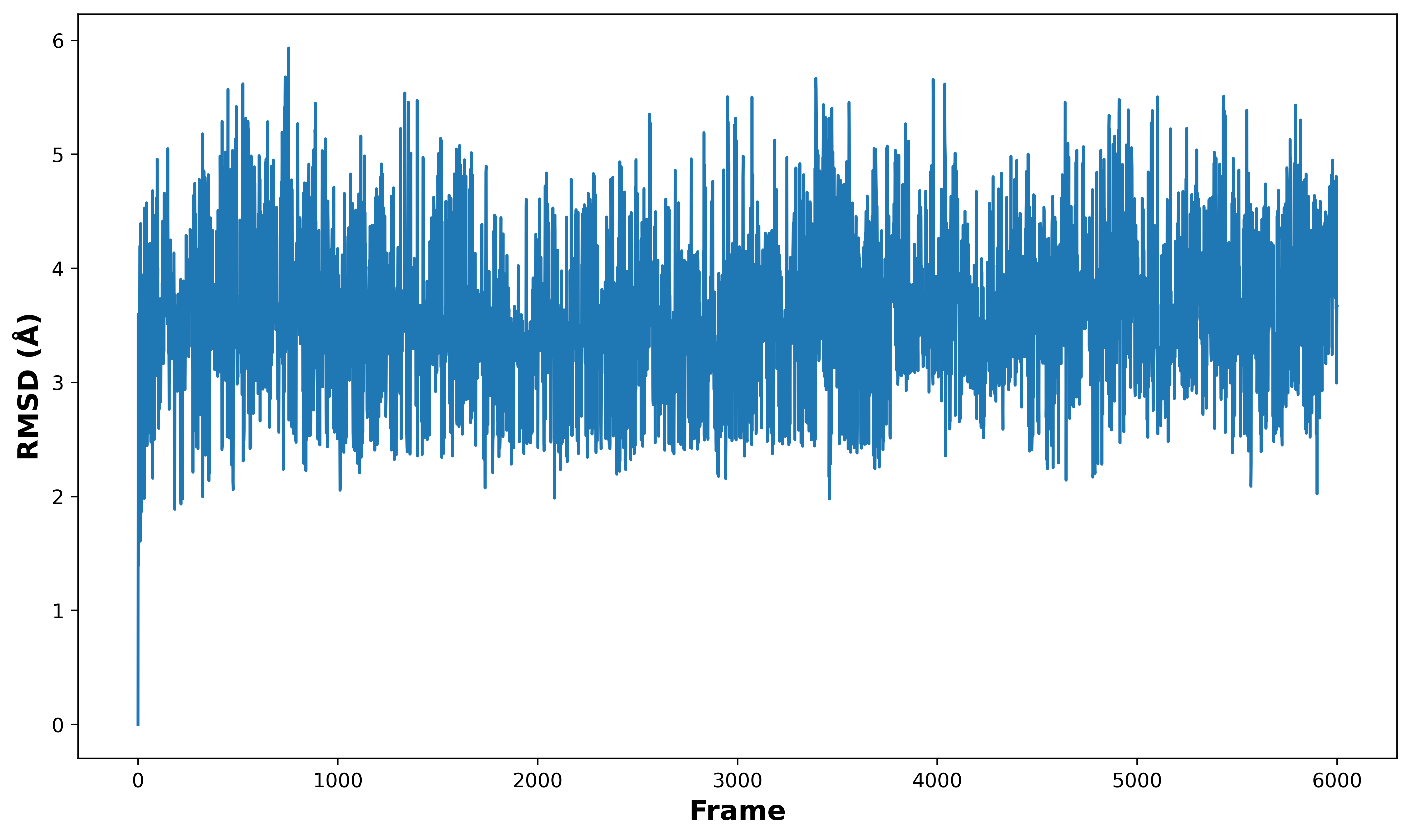

**Figure S6.** RMSD to the first frame for the β-heptapeptide trajectory plotted as a function of frame index. This trace provides a trajectory-level view of conformational variability across the full ensemble.

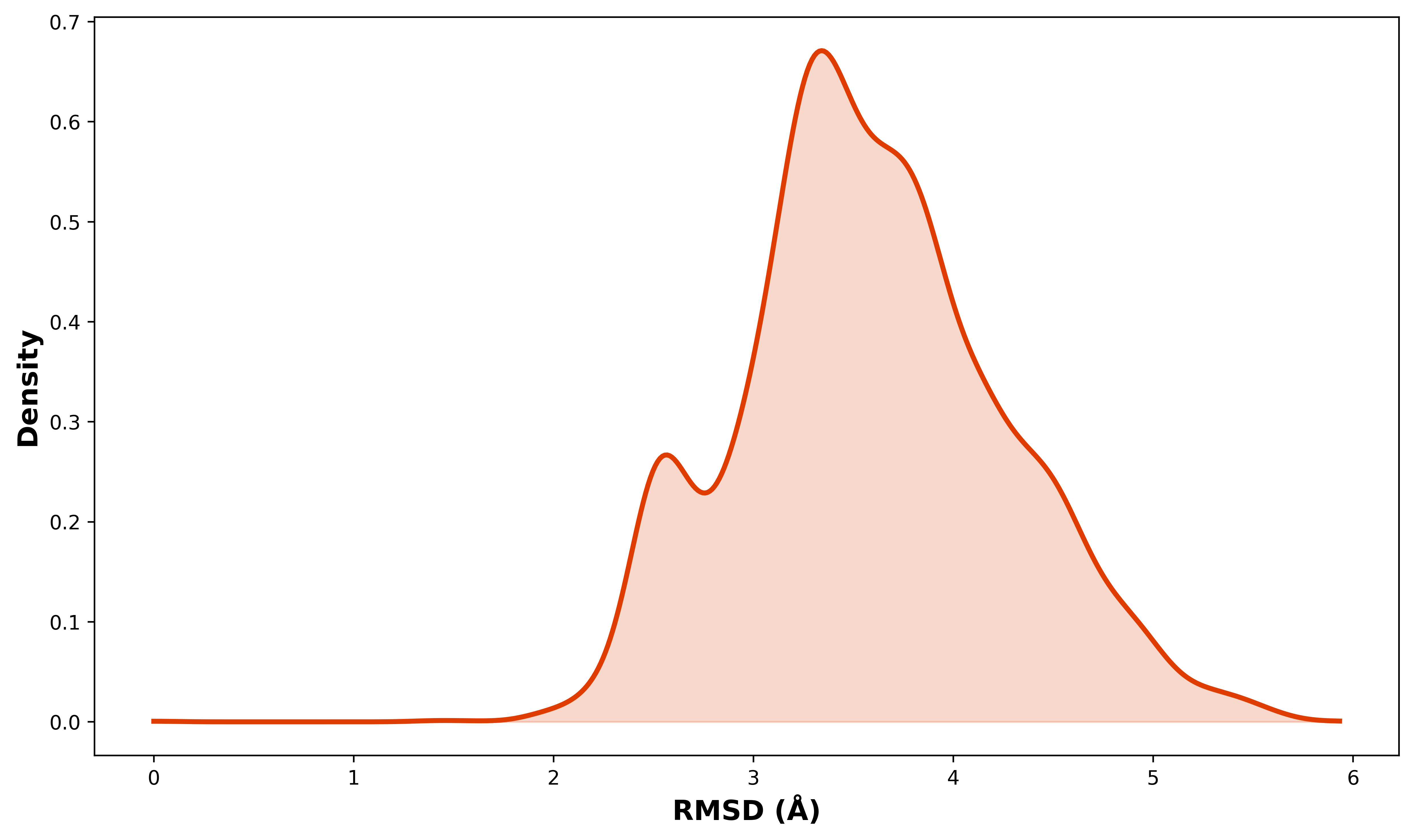

**Figure S7.** Distribution of RMSD to the first frame for the β-heptapeptide trajectory. The distribution summarizes the range of conformations sampled over the full trajectory.

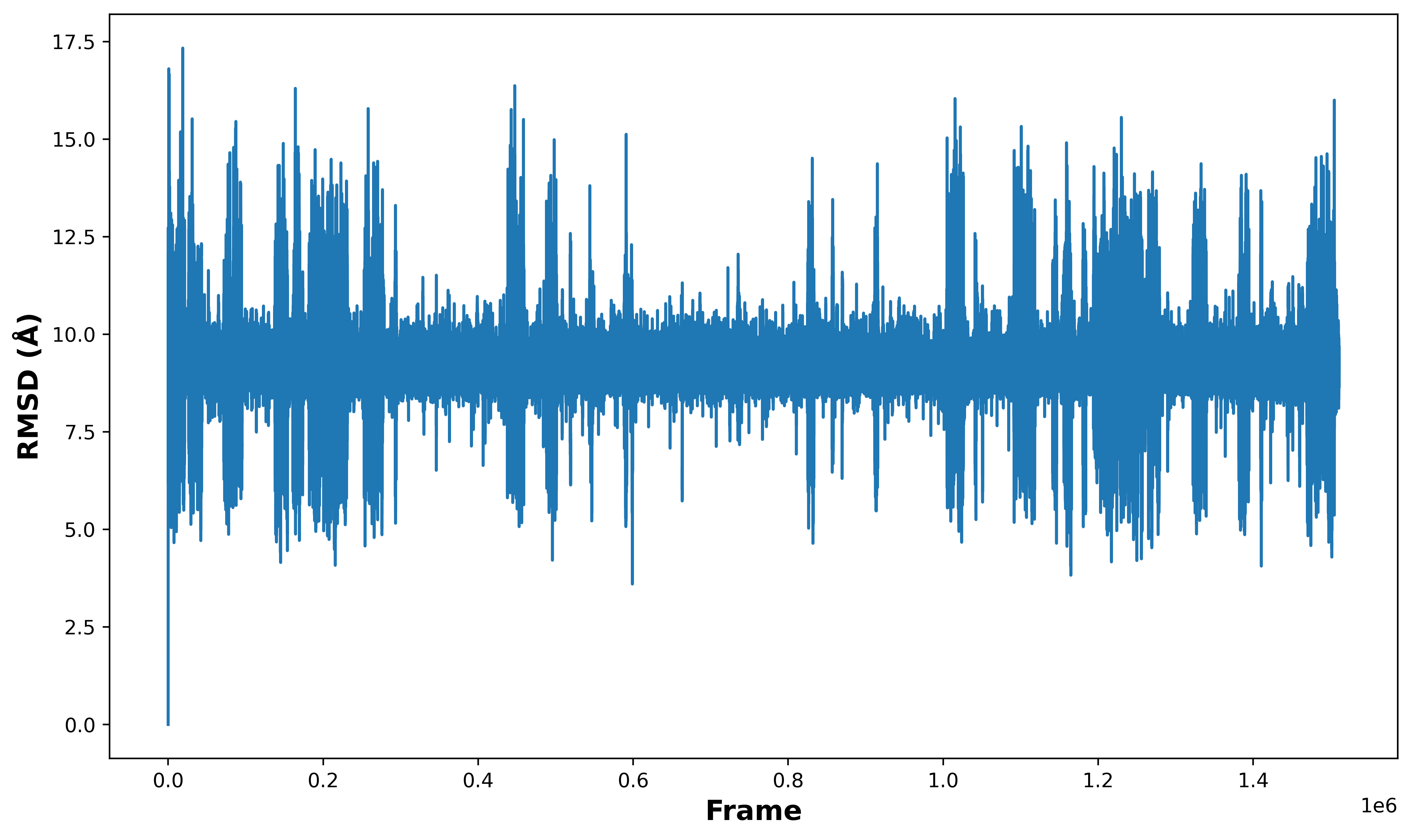

**Figure S8.** RMSD to the first frame for the HP35 trajectory plotted as a function of frame index. This trace provides a trajectory-level view of conformational variability across the full ensemble.

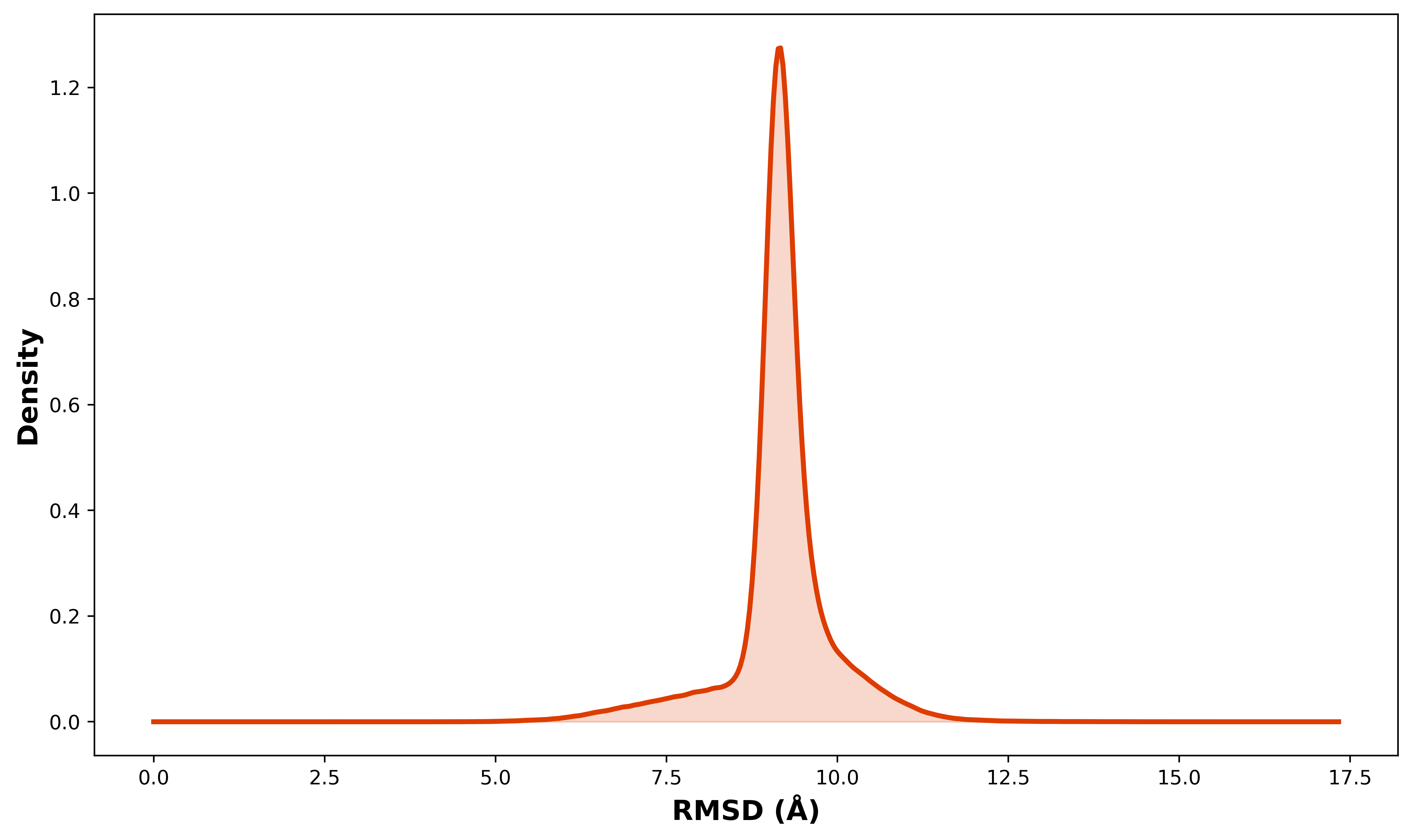

**Figure S9.** Distribution of RMSD to the first frame for the HP35 trajectory. The distribution summarizes the range of conformations sampled over the full trajectory.

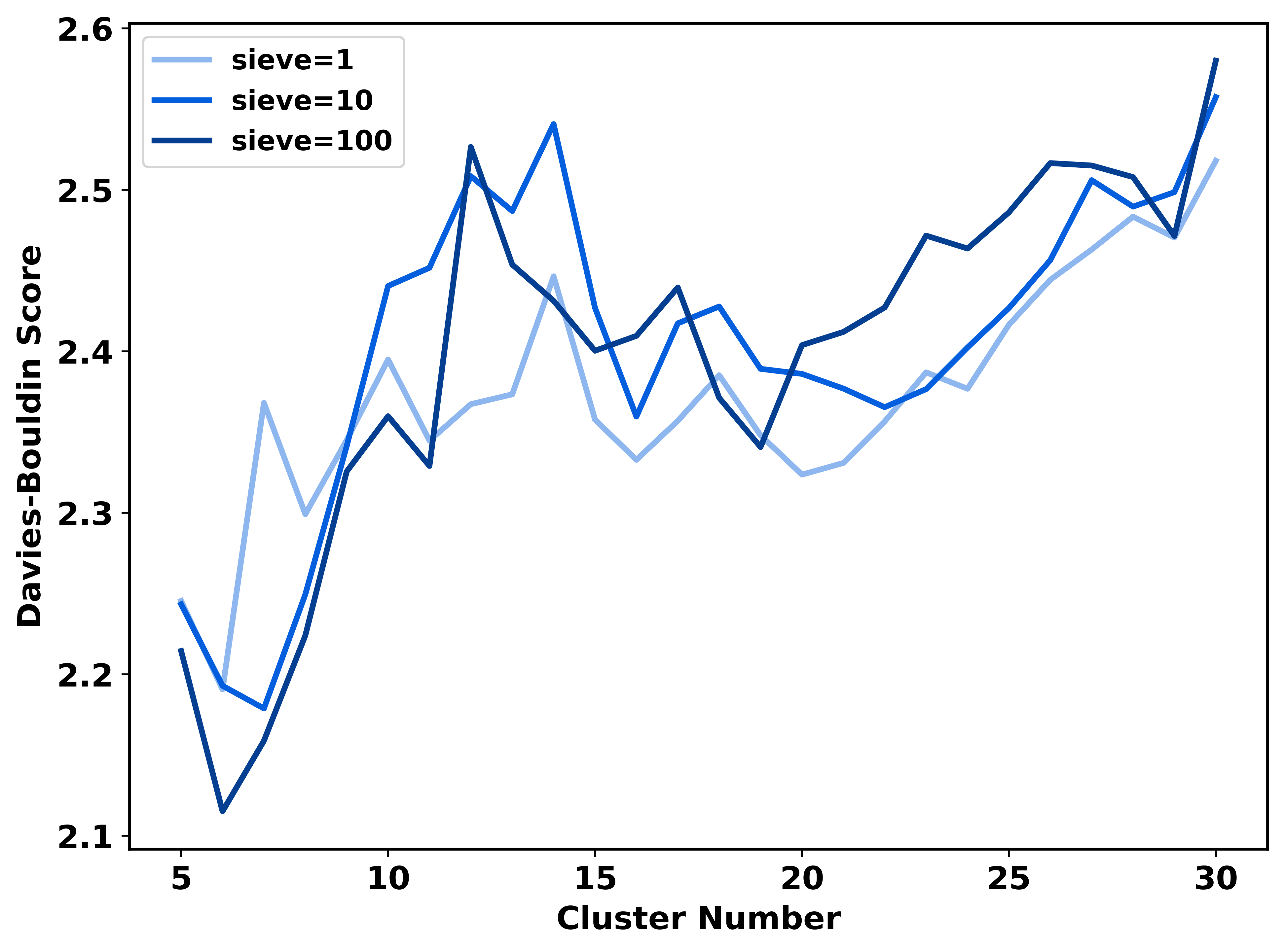

**Figure S10.** Davies-Bouldin index (DBI) for HP35 clustering at three sampling densities (sieve = 1, 10, 100) using comp_sim. DBI trends remain consistent across sieve values, indicating comparable clustering quality from the down-sampled sets to the full 1.5 million-frame trajectory.

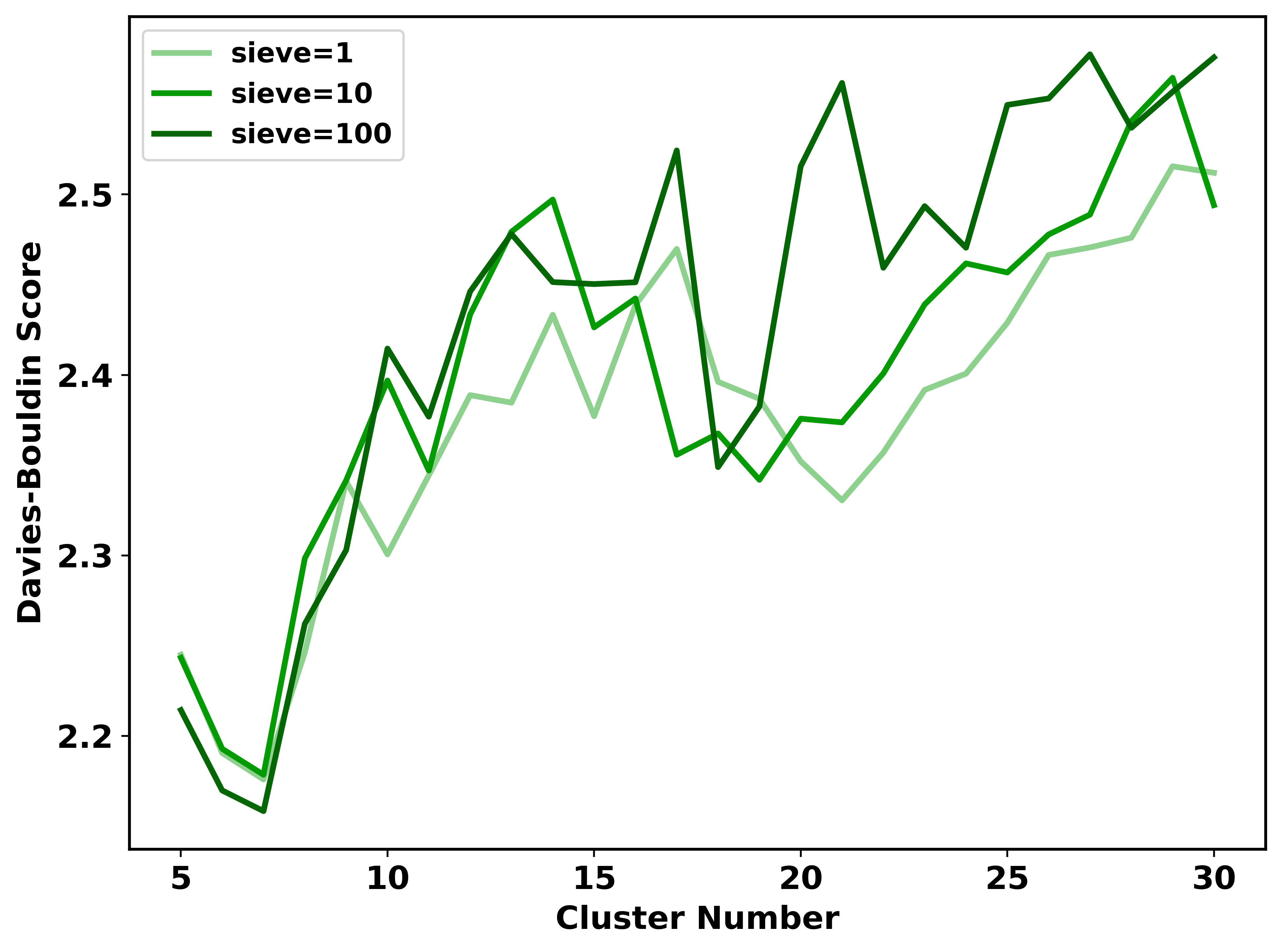

**Figure S11.** Davies-Bouldin index (DBI) for HP35 clustering at three sampling densities (sieve = 1, 10, 100) using strat_all. DBI trends remain consistent across sieve values, indicating comparable clustering quality from the down-sampled sets to the full 1.5 million-frame trajectory.

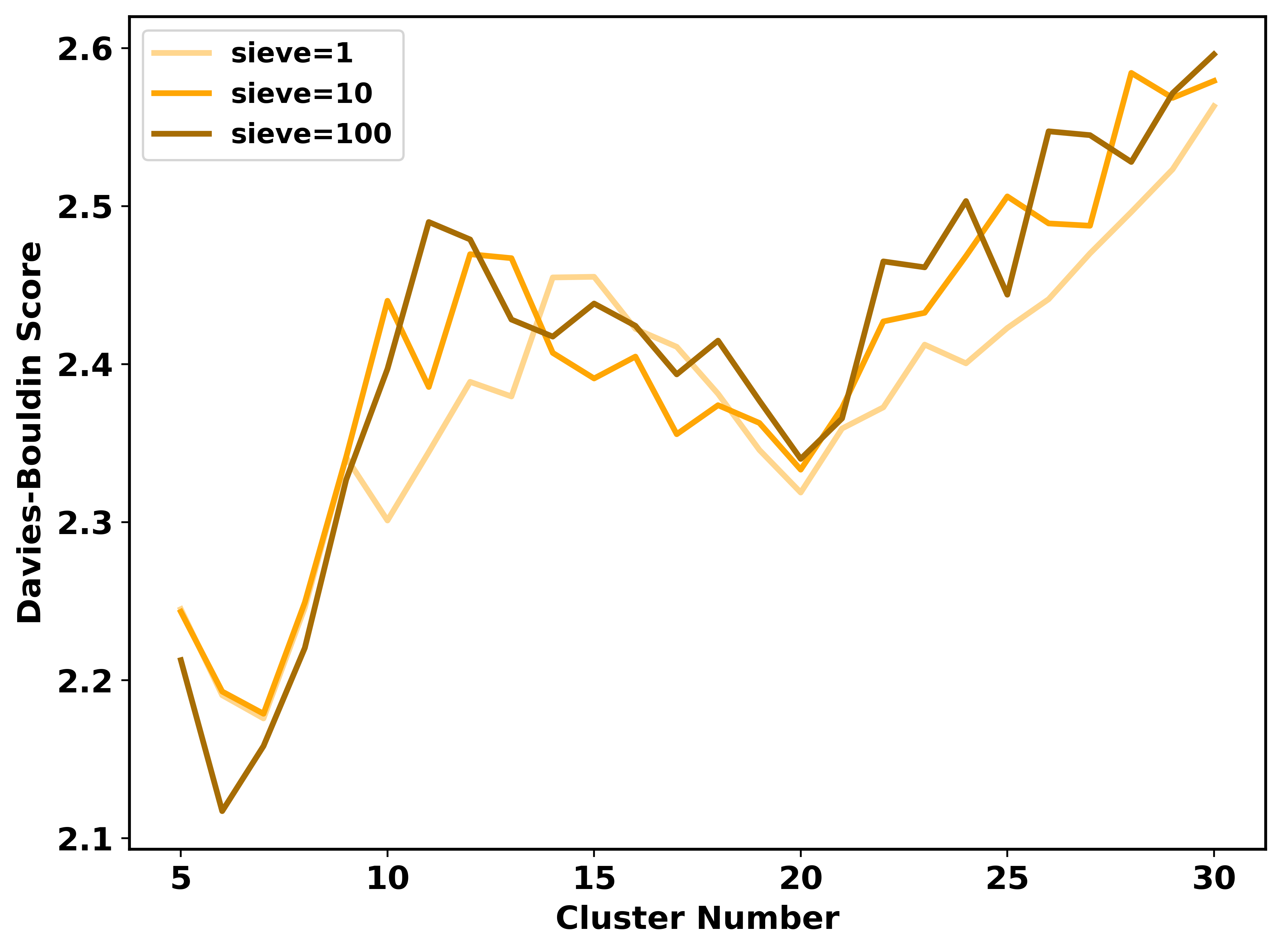

**Figure S12.** Davies-Bouldin index (DBI) for HP35 clustering at three sampling densities (sieve = 1, 10, 100) using strat_reduced. DBI trends remain consistent across sieve values, indicating comparable clustering quality from the down-sampled sets to the full 1.5 million-frame trajectory.
